# Unraveling *Xanthomonas* acetyltransferase GumG: decoding catalytic promiscuity, understanding the mechanism, and enhancing enzymatic versatility

**DOI:** 10.64898/2026.06.22.733816

**Authors:** Yajing Liu, Broder Rühmann, Okke Melse, Tenuun Bayaraa, Laura Kampl, Manuel Döring, Volker Sieber

**Affiliations:** Chair of Chemistry of Biogenic Resources, Technical University of Munich, TUM Campus Straubing for Biotechnology and Sustainability, Schulgasse 16, 94315 Straubing, Germany; Boehringer Ingelheim Pharma GmbH & Co. KG, Birkendorfer Str. 65, 88397 Biberach an der Riß, Germany; SynBioFoundry@TUM, Technical University of Munich, Schulgasse 22, 94315 Straubing, Germany; TUM Catalysis Research Center, Ernst-Otto-Fischer-Straße 1, 85748, Garching, Germany; School of Chemistry and Molecular Biosciences, The University of Queensland, 68 Copper Road, St. Lucia 4072, Australia

**Keywords:** Xanthan, Acetyltransferase GumG, Enzymatic modification, Enzyme engineering

## Abstract

Xanthan is a structurally complex exopolysaccharide produced by *Xanthomonas campestris* and one of the most extensively studied microbial biopolymers. As a sustainable alternative to petroleum-based polymers, its broader application requires precise control of polysaccharide decoration, yet the enzymatic basis of these modifications remains incompletely understood. Here, we characterise the activity and substrate scope of GumG, an AT-3 domain-containing membrane-bound acetyltransferase responsible for xanthan *O*-acetylation. Using mass spectrometry in combination with *in vitro* and *in vivo* assays, we show that GumG mediates non-specific acetylation of the outer mannose residue and displays pronounced substrate promiscuity. GumG also exhibits limited propionyltransferase activity, enabling the biosynthesis of hybrid acetylated-propionylated xanthan at an 8.27:1 ratio. Molecular docking and analysis of 31 xanthan variants identify a cytoplasmic substrate-binding pocket defined by Val67 and Phe71 that governs donor specificity, and an engineered GumG variant (F71L) shows enhanced propionyltransferase activity. In addition, a periplasmic His40-Trp143-Asp246-His297 motif is proposed to constitute the catalytic center. Together, these findings provide mechanistic insight into GumG multifunctionality and establish a framework for engineering xanthan derivatives with tailored physicochemical properties.

## 1. Introduction

The depletion of fossil-based resources and the escalating climate crisis have underscored the urgent need for sustainable, bio-based alternatives to petroleum-derived materials. Xanthan, a bacterial exopolysaccharide produced by *Xanthomonas campestris* (*Xcc*) with yields of up to 62 g L^−1^, has emerged as a promising candidate due to its biocompatibility, non-toxicity, and structural versatility [1]. The presence of functional groups in xanthan enables tailored modifications, allowing fine-tuning of its mechanical and biological properties. Although chemical synthesis of xanthan derivatives containing olefinic bonds has been explored for cross-linking applications, these approaches often suffer from limited specificity, inconsistent reproducibility, and undesirable environmental effects, motivating the development of biocatalytic strategies that provide precise and sustainable polymer modification under both *in vivo* and *in vitro* conditions [2, 3].

*O*-acetylation, a key modification in xanthan biosynthesis, is mediated by the acetyltransferases (ACTases) GumG and GumF [4], which belong to the Acyl_transf_3 (AT3) domain family. These integral membrane enzymes transfer acetyl groups from donors to specific acceptors—outer Man residues for GumG and inner Man residues for GumF [5–7]. AT3 domains, which typically contain ten transmembrane helices (TMHs), are widespread among bacteria and acetylate diverse cell surface polymers, including *O*-antigen, exopolysaccharides, and peptidoglycan, thereby modulating bacterial virulence and resistance [8–10]. AT3 proteins may function independently or as C-terminal SGNH-domain fusions (InterPro number IPR036514), with SGNH representing ∼200-aa GDSL hydrolase module; some carbohydrate-active SGNH members are classified as CE3 enzymes in CAZy [11], while OatA and OafB exemplify AT3–SGNH fusion architectures [9, 12]. Despite the central role of acetyl-CoA (Ac-CoA) in bacterial metabolism, its direct involvement in xanthan acetylation remains unresolved. Molecular dynamics simulations support a membrane-spanning Ac-CoA model that positions the acetyl group near extracytoplasmic catalytic residues [12]. Although the catalytic mechanism of AT3 enzymes is not fully elucidated, evidence from related ACTases suggests a role for catalytic triads or substrate-assisted catalysis [9, 13–15]. Two conserved AT3 motifs—an R/K-X_10_-H motif in TMH1 and a cytoplasmic RXXR motif in TMH3—are essential for activity, indicating a conserved mechanism for acetyl group positioning and transfer [12, 16].

Native xanthan adopts a double-helical conformation, and its rheological properties are strongly influenced by the distribution and degree of acetyl and pyruvyl substitution on side chains [5, 17, 18]. Strategies to modulate xanthan rheology have focused on engineering the *gum* gene cluster, notably *GumG*, *GumF*, and *GumL* [5, 7]. However, the enzymatic mechanisms underlying these modifications—particularly the balance between promiscuity and specificity—remain poorly understood, and enzyme engineering approaches to tune substrate selectivity are largely unexplored. Elucidation of these mechanisms underpins the rational design of functional biopolymers with tunable properties, including solubility, hydrophobicity and UV curability.

Here, comprehensive biochemical characterisation of GumG, an AT3-family ACTase from *Xanthomonas campestris*, reveals broad catalytic promiscuity that challenges the prevailing view of exclusive *O*-6 acetylation of outer mannose residues in xanthan. The enzyme accommodates diverse acyl-CoA donors—including Ac-CoA, propionyl-CoA (PA-CoA), and acryloyl-CoA (AA-CoA)—and modifies a range of acceptors, from monosaccharides to pentasaccharides, at multiple positions, thereby modulating xanthan hydrophobicity. Structural modeling and molecular docking suggest a mechanism for Ac-CoA translocation across the membrane and identify a catalytic center comprising His40, Trp143, Asp246, and His297 residues essential for pentasaccharide acetylation. Guided by these insights, an engineered GumG variant (F71L) exhibits enhanced propionyltransferase activity, illustrating the potential of rational enzyme design to functionalize biopolymers. Collectively, these findings expand the functional repertoire of AT3 ACTases and provide a framework for the sustainable production of tailored biopolymers with tunable properties.

## 2. Materials and Methods

### 2.1. Gene cloning and protein expression in *E. coli*

The truncated form of the xanthanase-encoding gene *RuGH5a*, which includes an N-terminal signal peptide, a catalytic GH5 domain, three carbohydrate-binding modules (CBM A, B, and C), and two *Listeria*-*Bacteroides* domains, as previously described by Ostrowski et al., was synthesized commercially (GeneArt Gene Synthesis, Thermo Fisher Scientific) to enable efficient pentasaccharide production [19]. The *Stlig* gene, encoding a 3-hydroxypropionyl-CoA synthetase, was amplified from chromosomal DNA of *Sulfolobus tokodaii* DSMZ 16993 via PCR for the synthesis of various CoA thioesters [20]. The *GumG* gene, encoding an ACTase, was PCR-amplified from *Xanthomonas campestris pv. campestris* LMG 8031 genomic DNA. For all genes, primer pairs were designed to incorporate in-frame 3’-extensions encoding a hexahistidine tag at the C-terminus of the target open reading frame (**Supplementary Table 1**). The resulting amplicons were cloned into the NcoI/XhoI-linearized *pET28* vector using Gibson Assembly Master Mix (New England Biolabs). *E. coli* DH5α competent cells were transformed with recombinant plasmids via heat shock transformation. Plasmid construction was verified by Sanger sequencing (Eurofins Genomics) before introduction into expression strains: *E. coli* BL21 DE3 for RuGH5a and Stlig expression, and *E. coli* C43 for GumG expression. Protein expression and purification in *E. coli* BL21 followed the previously described protocol [21], while expression and purification of GumG in *E. coli* C43 were performed as described by Jones *et al.* [9], with minor modifications. Briefly, cells harbouring the GumG-containing plasmid were grown to mid-log phase (OD_600_=0.5) and induced with 0.5 mM IPTG. After induction, the culture was incubated for an additional 6 h at 37°C and then harvested by centrifugation (10,000 g, 4°C, 20 min). Harvested cells were lysed by ultrasonication in 50 mM phosphate buffer (pH 7.0) containing 200 mM NaCl, 50 µg mL^−1^ lysozyme, 20 µg mL^−1^ DNase, 20 µg mL^−1^ RNase, and protease inhibitors (Roche). Unbroken cells were removed by centrifugation (11,000 g, 4°C, 20 min). The membrane fraction was isolated by ultracentrifugation (142,000 g, 4°C, 1 h) and then solubilized with DDM at 4 °C overnight, and GumG was subsequently purified by nickel-affinity chromatography.

### 2.2. Fermentative cultivation of *Xcc* (variants) and production of xanthan

The WT *Xcc* strain and its derivatives were cultured at 30 °C in MM1 medium (P100) supplemented with 10 g L⁻¹ Glc and 10 g L⁻¹ MOPS (pH 7.0) for xanthan production. The isopropyl-β-D-thiogalactopyranosid (IPTG)-inducible plasmid *pSRKGm* was used to express a C-terminal hexahistidine-tagged GumG ACTase and its variants in the *Xcc* Δ*gumFGL* deletion mutant, which was previously constructed by Gansbiller *et al.* to investigate the *in vivo* function of GumG [5]. Expression of GumG in the membrane fraction of *Xcc* Δ*gumFGL* was verified by Western blotting using anti-His-tag antibodies **(Supplementary Method 1, Supplementary Fig.1)**.

### 2.3. Pentasaccharides production, identification, and purification

Pentameric xanthan oligosaccharides with defined acetylation and pyruvylation patterns were generated by digesting xanthan variants (ΔFGLXan, ΔLXan, or ΔGLXan gum, produced by genetically modified *Xcc* strains—*Xcc* Δ*gumFGL*, *Xcc ΔgumL*, and *Xcc ΔgumGL*, respectively) with xanthanase RuGH5a [5, 19]. Briefly, 5 mg mL⁻¹ of each xanthan variant was incubated with 1.05 µM RuGH5a in 50 mM PBS (pH 7.0) at 37 °C for 3 days. The reaction mixture was precipitated with twice the volume of isopropanol, centrifuged, and the oligomer-containing aqueous phase was evaporated using a SpeedVac. The resulting powder was resuspended in 10 mL ultrapure water, centrifuged at 12,000 g for 5 min, and filtered through a 0.45 µm cellulose filter. The filtrate was purified using an ÄKTA pure 25L1 FPLC system (GE Healthcare) equipped with a 450 mL Bio-Gel P2 column, with water as the mobile phase [22]. Fractions containing reducing ends were first identified using the 3,5-dinitrosalicylic acid (DNS) assay, then derivatised with 1-phenyl-3-methyl-5-pyrazolone (PMP) and analyzed by LC-MS/MS [23, 24]. Fractions containing pentasaccharides were further pooled and concentrated using a SpeedVac for further enzyme assay.

### 2.4. Outer mannose of xanthan/pentasaccharides analysis

To determine the acetylation function of GumG during xanthan biosynthesis, the outer Man residues of the xanthan side chain were released by enzymatic digestion of xanthan polymer chains using xanthan lyase (X-L) [25]. Briefly, 101.54 nM of X-L enzyme was incubated with a 5 mg mL⁻¹ xanthan solution in a total reaction volume of 2 mL at 37 °C for 3 days. After digestion, the undigested or partially digested polymer was precipitated by adding two volumes of isopropanol. The supernatant, containing the liberated Man residues, was collected and concentrated using a SpeedVac. The dried material was resuspended in 200 µL of water and analyzed by LC-MS/MS.

### 2.5. *In vitro* production and determination of CoA thioesters

Different CoA thioesters, including Ac-CoA, AA-CoA, PA-CoA, crotonyl-CoA (CA-CoA), and methacryloyl-CoA (MAA-CoA), were synthesized using Stlig [20]. The reaction mixture contained 20 mM carboxylic acid (pH adjusted to 7.0 with NaOH), 0.25 mM CoA, 3 mM ATP, 10 mM MgCl₂, 100 mM Tris-HCl (pH 7.0), and 271.75 µM recombinant Stlig. Reactions were incubated at 37 °C for 10 min and quenched by freezing at −20 °C.

### 2.6. Analysis of GumG native activity and catalytic promiscuity

*In vitro* assay: The native activity of GumG was assessed using Ac-CoA as the donor and pentasaccharides as acceptors. Catalytic promiscuity was further evaluated through a standard transferase assay using various acyl donors (Ac-CoA, PA-CoA, AA-CoA, CA-CoA, and MAA-CoA) and monosaccharide or disaccharide acceptors (L-Man, D-Man, D-Glc, D-Gal, D-GlcA, and Man_2_). One-pot enzyme reactions were performed by incubating 7.85 µM GumG and 3.63 µM Stlig with 250 µM CoA, 20 mM carboxylic acid (pH 7.0), 3 mM ATP, and 55.6 mM monosaccharides (or 29.21 mM Man_2_ or 13.67 mM PenΔFGL or 8.08 mM PenΔGL) in 50 mM phosphate buffer (pH 7.0) containing 200 mM NaCl and 10 mM MgCl₂ at 30 °C, with 600 rpm shaking for two days. Negative controls were conducted by replacing GumG with buffer. Reaction products were derivatised with PMP and analysed by LC-MS/MS.

*In vivo:* To evaluate the *in vivo* substrate promiscuity of GumG, the *Stlig* gene was cloned into the IPTG-inducible vector *pSRKGm* and introduced into the *Xcc ΔgumL* (constructed by Ganzbiller *et al.* [5]) via electroporation, resulting in *Xcc ΔgumL* (*pStlig*). One of the following short-chain fatty acids (SCFAs; pH adjusted to 7.0), including 10 mM propionic acid, 10 mM methacrylic acid, 10 mM crotonic acid or 5 mM acrylic acid, was added to the culture medium of the *Xcc ΔgumL* (*pStlig*) strain concomitantly with induction by 1 mM IPTG at OD_600_=2.0-2.5. Preliminary experiments were conducted to determine the maximum acid concentration compatible with cell growth. After three days of cultivation, the polymers were isolated by precipitation with isopropanol, dried, and subsequently washed with water. The entire precipitation-drying-washing process was repeated three times. The purified polymers were either hydrolysed with 250 mM H_2_SO_4_ at 90 °C for 16 h and analyzed by high-performance liquid chromatography (HPLC), or digested with X-L and analyzed by LC-MS/MS.

### 2.7. Site-directed mutagenesis and plasmid construction

GumG variants with site-specific amino acid substitutions were generated using site-directed mutagenesis. Single amino acid replacements were introduced via PCR-based mutagenesis, followed by either T4 DNA ligation or Gibson assembly (**Supplementary Method 2**). The *pSRKGm*-*GumG* plasmid was employed for *in vivo* expression of GumG and its mutants in *Xcc* Δ*gumFGL*, where GumG catalyses xanthan modification. In contrast, *pET28*-*GumG* was used for heterologous expression in *E. coli* C43 to facilitate subsequent *in vitro* enzymatic assays. Primers designed for specific amino acid substitutions were used for PCR amplification **(Supplementary Table 1)**.

### 2.8. Bioinformatic analysis and site-directed mutagenesis of GumG

The 3D structure of GumG, predicted from its amino acid sequence, was generated using AlphaFold2. Surface electrostatic potential analysis was performed using APBS (Adaptive Poisson-Boltzmann Solver) implemented in PyMOL (https://pymolwiki.org/index.php/APBS). Potential binding sites for Ac-CoA in GumG were identified using AutoSite (https://ccsb.scripps.edu/autosite/). The conservation of all amino acid residues in GumG was analyzed using ConSurf (https://consurf.tau.ac.il/consurf_index.php) (**Supplementary Table 2**). Molecular docking simulations for Ac-CoA and the pseudo-xanthan pentameric unit were conducted using AutoDock Vina (https://vina.scripps.edu/). The docking strategy involved an initial docking of Ac-CoA, and the most favorable Ac-CoA-GumG complex was selected for subsequent docking with the pseudo-xanthan pentameric unit.

### 2.9. Metabolite quenching and extraction of intracellular CoA and CoA thioesters of *Xcc*

Metabolites were quenched using a cold acetonitrile:methanol:water (40:40:20) mixture. Samples were centrifuged, evaporated, and analyzed by LC-MS/MS (**Supplementary Method 3**).

### 2.10. Analytical methods

#### 2.10.1. Mono-/Oligomer composition of xanthan

Monosaccharides and pentasaccharides produced by enzymatic digestion were derivatised using the 1-phenyl-3-methyl-5-pyrazolone (PMP) method. Briefly, 25 µL of a 1 g L⁻¹ sugar solution was mixed with 75 µL of PMP master mix (125 mg PMP, 7 mL methanol, 3.0625 mL water, 437.5 µL 3.2% NH_4_OH) and incubated at 70 °C for 100 min. After derivatization, 20 µL of the sample was neutralized with 16.67 mM acetic acid in a final volume of 150 µL, centrifuged at 1,000 × g for 5 min, and then filtered through a 0.2 µm filter plate with a Supor® membrane prior to LC-MS/MS analysis.

Chromatographic separation for LC–MS/MS analysis was performed on a Gravity C18 column (100 mm × 2 mm, 1.8 µm; Macherey-Nagel) at 50 °C, using an UltiMate 3000RS HPLC system (Thermo Scientific, USA) coupled to an electrospray ionization ion-trap mass spectrometer (HCT, Bruker). Samples were maintained at 10 °C in the autosampler, and 10 µL was injected for each run. Data acquisition and analysis were conducted using Bruker Hystar, QuantAnalysis, and Dionex Chromeleon software. The mobile phase consisted of solvent A (5 mM ammonium acetate buffer, pH 5.6, containing 15% acetonitrile) and solvent B (acetonitrile). The gradient elution program was as follows: 0–5 min, 1–5% B; 5–7 min, 5% B; 7–8 min, 5–18% B; 8–8.3 min, 18–40% B; 8.3–10.3 min, 40% B; 10.3–10.5 min, 1% B; 10.5–12 min, 0% B. The flow rate was 0.6 mL min⁻¹, and PMP-labeled sugars were monitored at 245 nm. Prior to electrospray ionization mass spectrometry (ESI-MS), the LC effluent was split at a ratio of 1:20 using an Accurate-Post-Column-Splitter (Dionex). The mass spectrometer was operated in ultra-scan mode (scan rate: 26,000 m/z/s) over an m/z range of 50–1500. Source parameters were set as follows: nebulizer pressure: 40 psi; capillary voltage: 4 kV; skimmer voltage: 40 V; drying gas temperature: 325 °C; and drying gas flow: 6 L min^−1^. Data acquisition was performed in Auto-MS mode, using a smart target mass of 750 m/z and an MS/MS fragmentation amplitude of 0.5 V. The quantification was performed by using the extracted ion chromatograms (EIC) of the m/z value corresponding to the protonated molecules.

#### 2.10.2. Acetate, pyruvate and propionate Content

Xanthan was hydrolysed with 250 mM H₂SO₄ at 90 °C for 16 h [5]. After hydrolysis, samples were cooled to room temperature, centrifuged at 14,000 rpm for 1 min, and filtered through a 0.22 µm PVDF syringe filter. Acetate, pyruvate, acrylate, and propionate were quantified using an Ultimate-3000 HPLC system equipped with a Rezex™ ROA-Organic Acid H⁺ column (Phenomenex). Chromatographic separation was carried out at 70 °C with a flow rate of 0.5 mL min⁻¹ for 35 min, using 2.5 mM H₂SO₄ as the mobile phase. Compound concentrations were determined from standard curves (0.5–10 mM) prepared with sodium acetate, sodium pyruvate, acrylate, and propionate, and detected using a UV detector and a refractive index (RI) detector.

#### 2.10.3. CoA thioester analysis

Analysis of CoA thioesters was performed using a porous reversed-phase column (Luna® Omega Polar C18, 1.6 µm, 100 mm × 2.1 mm, Aschaffenburg, Germany). Samples were maintained at 10 °C in the autosampler, and 10 µL was injected for each analysis. Chromatographic separation was performed at 50 °C with a flow rate of 0.6 mL min^−1^, using 26.5 mM ammonium formate (pH 6.5, adjusted with 3.2 % ammonia) as the aqueous mobile phase and a gradient elution of acetonitrile over 13 minutes, as follows: 0-2 min: 0 % acetonitrile, 2-7 min: linear gradient from 0 to 50% acetonitrile, 7-9 min: 50 % acetonitrile, 9-9.1 min 50-0 % acetonitrile, 9.1-13 min: re-equilibration at 0 % acetonitrile. CoA thioesters were identified and quantified based on their specific mass-to-charge (m/z) transitions, involving characteristic parent and fragment ions. The mass spectrometer settings were the same as those used for the PMP method.

#### 2.10.4. GDP-mannose analysis

Samples were diluted to a final concentration of 0.5 mM GDP-mannose (GDP-Man). A 300 µL diluted sample was mixed with 700 µL of acetonitrile, and the resulting mixture was filtered through a 0.25 µm DTEF membrane to remove particulate matter prior to analysis. Chromatographic separation of GDP-Man was carried out using a hydrophilic-interaction liquid chromatography (HILIC) column (Triart Diol HILIC, 1.9 µm, 100 mm x 2.0 mm). The mobile phases consisted of water containing 0.1% (v/v) formic acid (Solvent A, pH 4.5) and acetonitrile containing 0.1% formic acid (Solvent B). The flow rate was maintained at 0.4 mL min^−1^. Gradient elution of Solvent B was performed over 15 minutes as follows: 0–2 min, 85% B; 2–4 min, linear decrease from 85% to 79% B; 4–5 min, 79% B; 5–8 min, linear decrease from 79% to 70% B; 8–10 min, 70% B; and 10–15 min, re-equilibration at 85% B. The mass spectrometer settings were the same as those used for the PMP method. GDP-Man, both with and without acetylation, was identified and quantified based on its specific mass-to-charge (m/z) transitions.

## 3. Results

### 3.1. Native activity and catalytic promiscuity of GumG in xanthan *O*-acetylation

#### 3.1.1. *In vivo* assessment of GumG native activity

To investigate the *in vivo* role of GumG in xanthan modification, four structurally distinct xanthan variants were generated: wild-type xanthan (WT Xan) from *Xanthomonas campestris pv. campestris* LMG 8031 (*Xcc*; BCCM, Gent, Belgium); XanΔFGL, lacking both acetylation and pyruvylation; XanΔGL, deficient in GumG and GumL, and predicted to lack acetylation and pyruvylation of the outer Man; and XanΔFL, lacking GumF and GumL, and presumed to contain a non-acetylated inner Man and a non-pyruvylated outer Man. Quantification of acetate and pyruvate released after acid hydrolysis indicated mannose modification degrees of 100.0% for acetylation and 54.8% for pyruvylation in WT Xan. In contrast, XanΔFGL contained only trace amounts of both substituents **(Fig. 1a, b)**. Notably, partial acetylation was retained in XanΔFL and XanΔGL, consistent with compensatory activities of GumG and GumF, respectively, as reported previously [5]. However, HPLC-based compositional quantification cannot resolve site-specific acetylation.

**Fig. 1.**
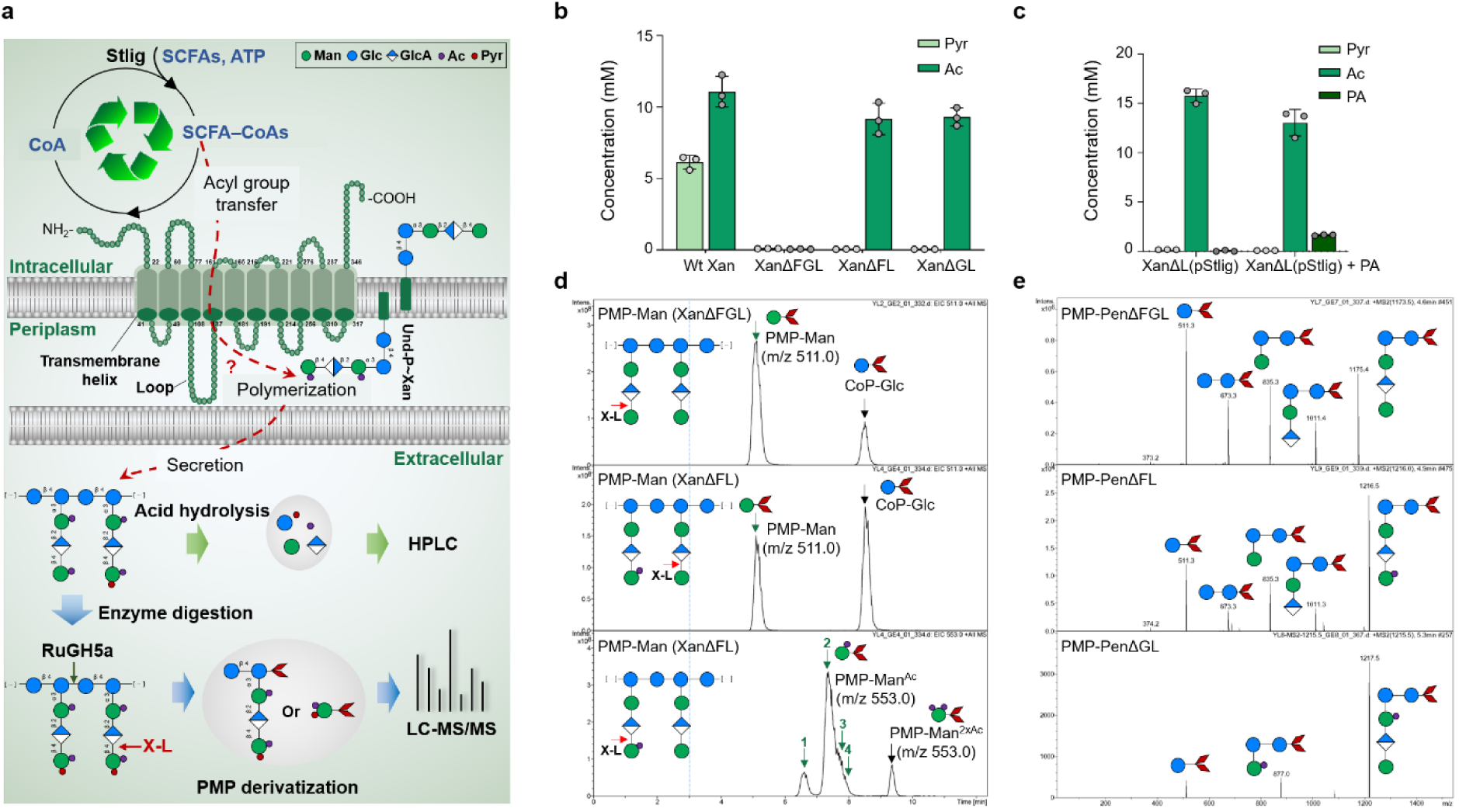
Structural and functional characterisation of GumG and *in vivo* xanthan modification. **a,** Predicted membrane topology and proposed function of *X. campestris* GumG (Membrain 3.1). GumG is hypothesised to catalyse acetylation at the outer Man residue of the xanthan pentameric repeat unit. The schematic illustrates the cyclic biocatalytic strategy for *in vivo* xanthan modification, in which Stlig generates SCFA–CoA thioesters, GumG transfers the acyl group to xanthan, and the released CoA is recycled. Analytical workflows for mono-/oligosaccharide and acyl-group analyses are indicated. **b, c,** Quantification of Ac, Pyr and PA in xanthan hydrolysates. Xanthan polymers were produced by wild-type (WT), engineered strains (*Xcc ΔgumFGL*, *Xcc ΔgumFL* and *Xcc ΔgumGL*), and Stlig-expressing *Xcc ΔgumL (pStlig)* cultured in the presence of 10 mM propionic acid. Polymers (10 g L⁻¹) were hydrolyzed in 250 mM H₂SO₄ at 90 °C for 16 h, followed by HPLC analysis using a Rezex™ ROA H⁺ column. **d,** Outer Man acetylation profiles of XanΔFGL and XanΔFL. Extracted ion chromatograms (EICs) show signals at m/z 511 corresponding to non-acetylated D-Man and at m/z 553 corresponding to acetylated D-Man^Ac^. Four distinct signals (green arrows) at m/z 553 indicate spatially non-specific GumG-mediated acetylation of outer Man residues in XanΔFL. A signal indicated by a black arrow corresponds to doubly acetylated outer mannose residues. This assignment is supported by a low-intensity signal at m/z 575 observed at the same retention time. The dominant signal at m/z 553 likely arises from in-source loss of one acetyl group during ionisation, reflecting the lability of the second acetyl moiety. **e,** MS^2^ spectra of xanthan pentameric units. Fragment ions correspond to PenΔFGL (m/z 1173.5) and to PenΔFL and PenΔGL (m/z 1215.5). Pentamers (generated using 1.05 µM RuGH5a) and free outer Man residues (generated using 101.54 nM X-L) were obtained by targeted enzymatic digestion of xanthan (5 g L⁻¹), followed by PMP derivatisation and LC-MS/MS analysis. **Symbol legend:** red rhombus, PMP; blue circle, glucose (Glc); green circle, mannose (Man); blue-and-white diamond, glucuronic acid (GlcA); purple circle, acetate (Ac); red circle, pyruvate (Pyr). **Abbreviations:** SCFA, short-chain fatty acid; SCFA–CoA, short-chain fatty acyl–coenzyme A; CoP-Glc, co-precipitated glucose.

To map acetylation sites, xanthan variants were enzymatically depolymerized using xanthan lyase (X-L), which cleaves off the terminal mannose residue independently of its decoration, or RuGH5a, and the resulting mono- and oligosaccharides were analyzed by LC-MS/MS (**Fig. 1a)**. Analysis of xanthan lyase digests from XanΔFL revealed both non-acetylated (m/z 511) and acetylated (m/z 553) PMP-derivatised mannose species, whereas the acetylated form was absent from XanΔFGL **(Fig. 1d)**. These data verified that GumG mediates acetylation of the terminal mannose residue, as described before. Surprisingly, the presence of four distinct chromatographic peaks corresponding to the acetylated species indicates heterogeneous acetylation, most likely at the *O*-2, *O*-3, *O*-4 and *O*-6 positions (with retention times increasing accordingly), thereby challenging the prevailing assumption that xanthan acetylation occurs exclusively at the *O*-6 acetylation [26].

Consistently, RuGH5a digestion of XanΔFGL yielded unmodified pentameric oligosaccharides (m/z 1173.5), whereas digests of XanΔFL and XanΔGL were dominated by monoacetylated pentamer (m/z 1215.5, + 42 Da) (**Fig. 1e**). MS/MS analysis of the m/z 1173.5 species yielded fragment ions consistent with an unmodified pentamer (**Fig. 1e**). A diagnostic fragment at m/z 1215.5 was observed exclusively in XanΔFL compared with XanΔFGL, supporting GumG-dependent acetylation of the outer mannose residue.

Additional analyses of WT Xan and XanΔGL, presented in the **Supplementary Fig. 2-4**, are consistent with GumF-mediated acetylation of the inner mannose residue and support a coordinated role of GumG, GumF and GumL in shaping the acetylation profile of native xanthan.

#### 3.1.2. *In vivo* assessment of GumG catalytic promiscuity

The substrate promiscuity of GumG was investigated using the *Xcc ΔgumL* strain, which encodes GumG and GumF, transformed with an IPTG-inducible *pSRKGm* plasmid carrying *Stlig* (designated *Xcc ΔgumL* (*pStlig*)). Stlig converts short-chain fatty acids into their corresponding CoA thioesters *in vitro,* was confirmed by LC-MS/MS analysis **(Supplementary Fig. 5-6)**. The *in vivo* catalytic promiscuity of GumG was examined by supplementing *Xcc ΔgumL* (*pStlig*) cultures with acrylic, propionic, crotonic, or methacrylic acids at the highest concentrations compatible with growth **(Supplementary Fig. 7)**, followed by purification of the resulting polymers to remove residual free acids. Acid hydrolysis with H_2_SO_4_ followed by HPLC analysis detected propionic acid at concentrations of 15.6 µM and 1.65 mM in polymers derived from acrylic- and propionic acid-supplemented cultures, respectively. These values corresponded to calculated propionate contents of 0.14% and 14.54%. In contrast, acrylic acid was not detected in the hydrolysates, potentially reflecting limitations of the in-house analytical method **(Fig. 1c, Supplementary Fig. 8)**. In the negative control polymer (without propionic acid supplementation), neither acrylic acid nor propionic acid was detected in the hydrolysates.

Enzymatic digestion of the propionic acid-derived polymer with X-L, followed by PMP derivatisation and LC-MS/MS, confirmed the presence of propionylated outer Man residues (m/z 567). MS^2^ analysis identified a propionyl group (m/z 56) esterified to the outer Man, indicating transfer of the propionyl group to xanthan by GumG **(Supplementary Fig. 9)**. In contrast, no incorporation of crotonyl or methacryloyl groups was detected. Together, these results demonstrate that GumG can accept certain selected short-chain acyl donors *in vivo*, revealing its substrate flexibility in polysaccharide modification.

### 3.2 *In vitro* assessment of GumG native activity and catalytic promiscuity

#### 3.2.1. GumG utilizes acetyl-CoA for pentameric xanthan repeating unit *O*-acetylation

Although direct evidence has been lacking, Ac-CoA has been proposed as a potential acetyl donor for xanthan acetylation [27]. To evaluate this possibility *in vitro*, the ACTase activity of WT GumG was examined using a transferase assay with Ac-CoA as the acetyl donor and two xanthan-derived pentasaccharides, PenΔFGL and PenΔGL, as acceptors. These substrates were generated by RuGH5a digestion of XanΔFGL and XanΔGL, respectively **(Fig. 2a)**. Reactions lacking GumG were included as negative controls.

**Fig. 2.**
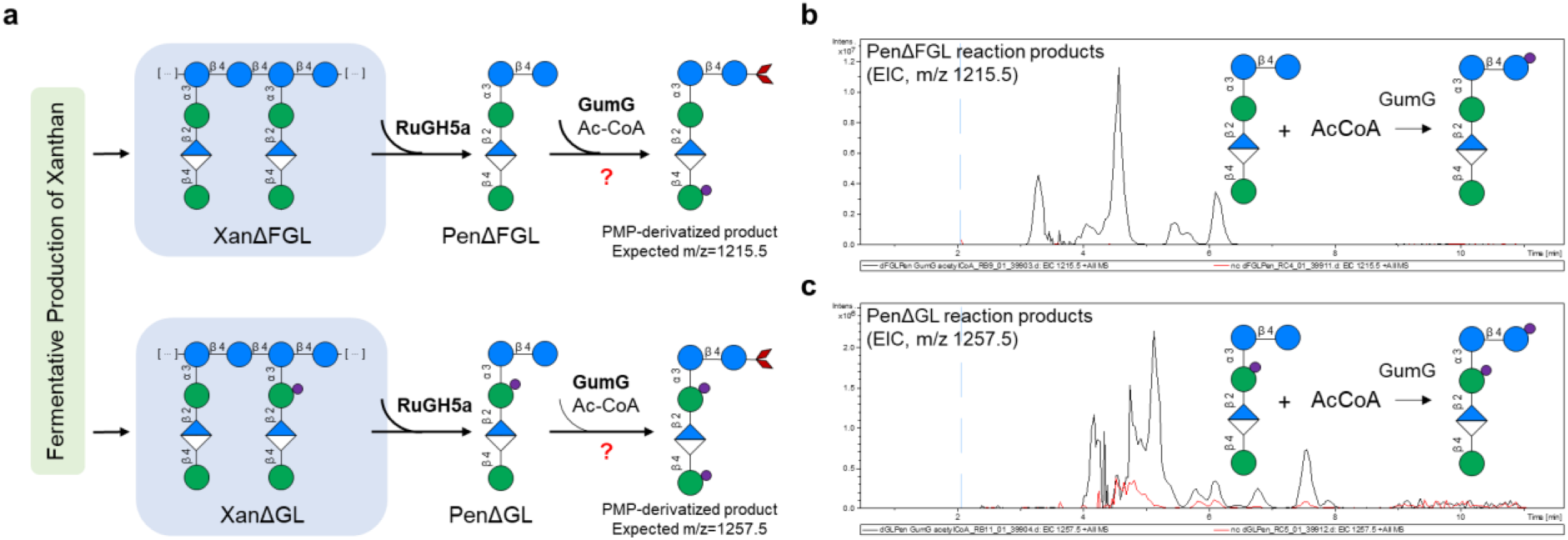
Catalytic promiscuity of GumG toward xanthan pentameric units. **a,** Schematic illustration of the generation of pseudo-pentameric xanthan repeat units by digestion with the xanthanase RuGH5a, followed by purification and use as acceptor substrates. **b,c,** Extracted ion chromatograms (EICs) of products from transferase assays using Ac-CoA as the donor and **b,** PenΔFGL or **c,** PenΔGL as acceptors. Reaction products formed in the presence of GumG are shown in black, whereas negative controls lacking GumG are shown in red. In **b,** the peak at m/z 1215.5 corresponds to singly acetylated PenΔFGL. In **c,** peaks at m/z 1257.5 represent GumG-catalysed acetylation of PenΔGL. Substrate and product schematics with corresponding m/z values are shown in **a**. Pentameric xanthan oligosaccharides were generated by treating xanthan (5 g L⁻¹) with 1.05 µM RuGH5a at 37 °C for 72 h. Purified pentasaccharides were incubated in transferase assays with 7.85 µM GumG, 100 mM Ac-CoA and either 13.67 mM PenΔFGL or 8.08 mM PenΔGL in 50 mM phosphate buffer (pH 7.0) at 30 °C and 600 rpm for 48 h. Negative controls were performed by replacing GumG with buffer. Reaction products were derivatised with PMP and analysed by LC-MS/MS. **Symbol legend:** red rhombus, PMP; blue circle, glucose (Glc); green circle, mannose (Man); blue-and-white diamond, glucuronic acid (GlcA); purple circle, acetate (Ac).

LC-MS/MS analysis revealed acetylated products when either PenΔFGL or PenΔGL was supplied as the acceptor, whereas no acetylation was detected in the absence of GumG. In reactions containing PenΔFGL, extracted ion chromatograms showed at least seven peaks at m/z 1215.5, corresponding to a +42 Da mass shift from the unmodified pentamer (m/z 1173.5), consistent with monoacetylation. The multiple peaks with identical precursor masses likely represent chromatographically resolved monoacetylated isomers, indicating that PenΔFGL generates structurally heterogeneous acetylated product (**Supplementary Fig. 10, Fig. 2b**).

In reactions using PenΔGL, partial acetylation of the inner Man residue by GumF generated a mixed population of acetylated and non-acetylated pentamers, while leaving the outer Man unmodified. Subsequent analysis focused on the acetylated inner-Man species (m/z 1215.5) to assess GumG activity. Additional ions at m/z 1257.5 were detected exclusively in reactions containing GumG, indicating further acetylation of PenΔGL **(Fig. 2c)**. MS/MS analysis of acetylated products from both PenΔFGL-and PenΔGL-based reactions unexpectedly localized the predominant acetylation site to the first glucose residue from the reducing end of the backbone, rather than to the outer Man **(Supplementary Fig. 10)**.

The signal intensity of GumG-mediated acetylated products was higher when PenΔFGL (m/z 1215.5) was used as the acceptor than when PenΔGL (m/z 1257.5) was used, despite the higher concentration of PenΔGL in the assay **(Fig. 2b,c)**, suggesting that GumG preferentially acts on fully non-acetylated pentasaccharide substrates.

#### 3.2.2. Catalytic promiscuity of GumG towards different acceptors

To evaluate the catalytic promiscuity of GumG as an *O*-ACTase, its activity was examined using a panel of small-molecule acceptors, including GDP-Man, monosaccharides (D-Man, D-Glc, D-Gal, L-Man and D-GlcA), and the disaccharide Man_2_.

When GDP-Man was used as the acceptor, LC-MS/MS analysis revealed a product with a mass of 647 Da, corresponding to monoacetylated GDP-Man (+42 Da relative to the native substrate), which was absent from reactions lacking GumG **(Supplementary Fig. 11)**. The acetylated product exhibited a shorter retention time than GDP-Man, consistent with increased hydrophobicity upon acetylation.

LC-MS/MS analysis further detected acetylated derivatives of D-Man, D-Glc, D-Gal and Man_2_, with characteristic ions at m/z 553 for the monosaccharides and m/z 715 for Man_2_ **(Fig. 3a)**. In contrast, negative controls contained only the corresponding non-acetylated species (m/z 511 for monosaccharides and m/z 673 for Man_2_). The acetylated sugars consistently exhibited longer retention times, and multiple chromatographic peaks were observed for each product, indicating acetylation at multiple hydroxyl positions and increased hydrophobicity. Quantitative comparison of signal intensities revealed substantially higher GumG activity toward D-Glc than toward D-Man or D-Gal, whereas no detectable acetylation was observed with L-Man or D-GlcA as acceptors **(Fig. 3a)**.

**Fig. 3.**
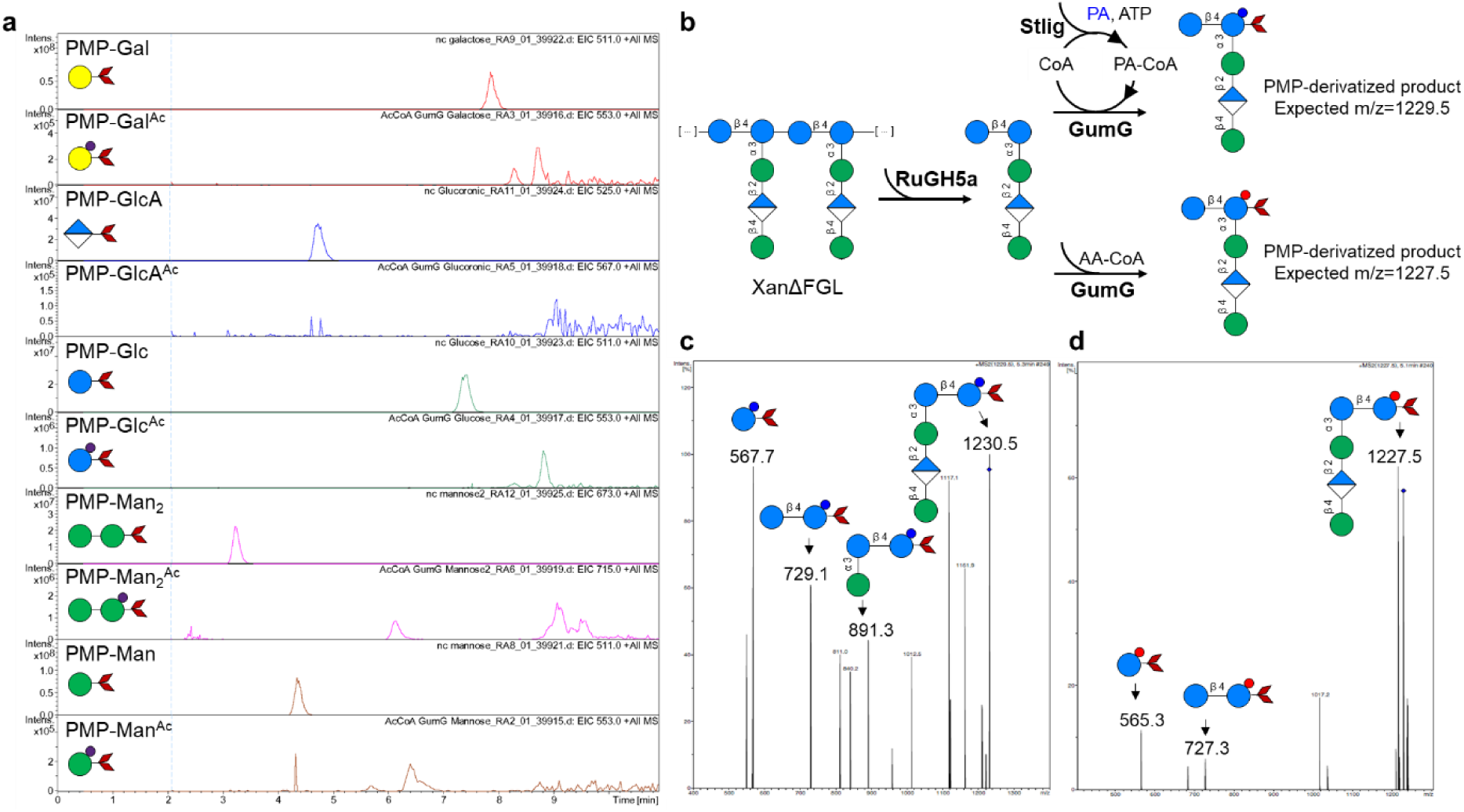
LC-MS/MS analysis of products from GumG-catalysed transferase assays. **a,** Extracted ion chromatograms (EICs) showing acetylated products detected exclusively in reactions containing GumG, an acyl donor and a sugar acceptor. Signals at *m/z* 553 correspond to PMP-derivatised acetylated D-galactose (D-Gal^Ac^), D-glucose (D-Glc^Ac^) and D-mannose (D-Man^Ac^). A peak at m/z 712 indicates PMP-derivatised acetylated mannobiose (Man ^Ac^). EICs of non-acetylated sugars show signals at m/z 511 for D-Gal, D-Glc and D-Man, at m/z 525 for D-glucuronic acid (D-GlcA), and at m/z 673 for mannobiose (Man2). Reactions were performed using acetyl-CoA (Ac-CoA) as the donor with 275 mM of each sugar acceptor and 8.25 µM GumG at 30 °C for 48 h; negative controls lacking GumG were conducted under identical conditions. **b,** Schematic illustration of pseudo-pentameric xanthan repeat units generated by digestion with the xanthanase RuGH5a, purified and used as acceptors, and PA-CoA generated *in situ* by Stlig and employed as the acyl donor. Owing to the intrinsic instability of AA-CoA, commercially available AA-CoA was used in acrylation assays. **c,** MS^2^ fragmentation of propionylated PenΔFGL, yielding diagnostic ions at m/z 567.7, 729.1, 891.3 and 1229.5, consistent with transfer of a propionyl group (PA) to the reducing-end Glc residue of the backbone. **d,** MS^2^ fragmentation of acrylated PenΔFGL, yielding ions at m/z 565.3, 727.3 and 1227.5, indicating transfer of an acryloyl group (AA) to the reducing-end Glc residue. One-pot transferase assays were conducted under two conditions. For propionyl transfer, GumG and Stlig were incubated with CoA, propionic acid, ATP and PenΔFGL. For acrylation, AA-CoA was incubated with PenΔFGL and GumG. Reaction products were analysed by LC-MS/MS. **Symbol legend:** red rhombus, PMP; yellow circle, galactose (Gal); blue circle, glucose (Glc); green circle, mannose (Man); blue-and-white diamond, glucuronic acid (GlcA); dark blue circle, propionyl group (PA); red circle, acryloyl group (AA).

#### 3.2.3. Catalytic promiscuity of GumG towards different donors

To assess the catalytic promiscuity of GumG, a cyclic biocatalytic system for sugar modification was established by coupling Stlig-mediated CoA thioester formation with GumG-catalysed acyl transfer in a one-pot reaction format. In this system, acyl groups generated *in situ* as CoA thioesters were transferred to sugar acceptors by GumG, while the released CoA was recycled to sustain continuous turnover (**Fig. 3b**).

Using PenΔFGL as the acceptor, LC–MS/MS analysis of reactions containing GumG, Stlig and propionic acid identified a product at m/z 1229.5, corresponding to a +56 Da mass shift relative to unmodified PenΔFGL (m/z 1173.5), consistent with propionylation. MS/MS fragmentation yielded diagnostic ions at m/z 567.7 (propionylated hexose), 729.1 (propionylated disaccharide) and 891.1 (propionylated trisaccharide), localizing the modification to the reducing-end glucose residue of the backbone (**Fig. 3c**).

Owing to the intrinsic instability of AA-CoA, reactions with acrylic acid were performed using 0.5 mM commercially available AA-CoA. LC–MS/MS analysis detected a product at m/z 1227.5, consistent with acrylation of the pentamer (+54 Da). Fragment ions at m/z 565 and 727 further localized attachment of the acryloyl group to the reducing-end glucose residue **(Fig. 3d)**. By contrast, no acyl transfer was detected when crotonic acid or methacrylic acid was supplied, consistent with the *in vivo* substrate specificity observed for GumG.

### 3.3. *In silico* prediction of the acetyl-CoA binding site in GumG

In the absence of experimentally resolved GumG structures, AlphaFold2 predictions combined with AutoDock docking were used to explore potential Ac-CoA and pentasaccharide binding sites. Together with Consurf-based conservation analysis of 46 homologues, these analyses provide mechanistic insights and a structural basis for engineering GumG‘s catalytic versatility (**Fig. 4a, Supplementary Table 2**).

**Fig. 4.**
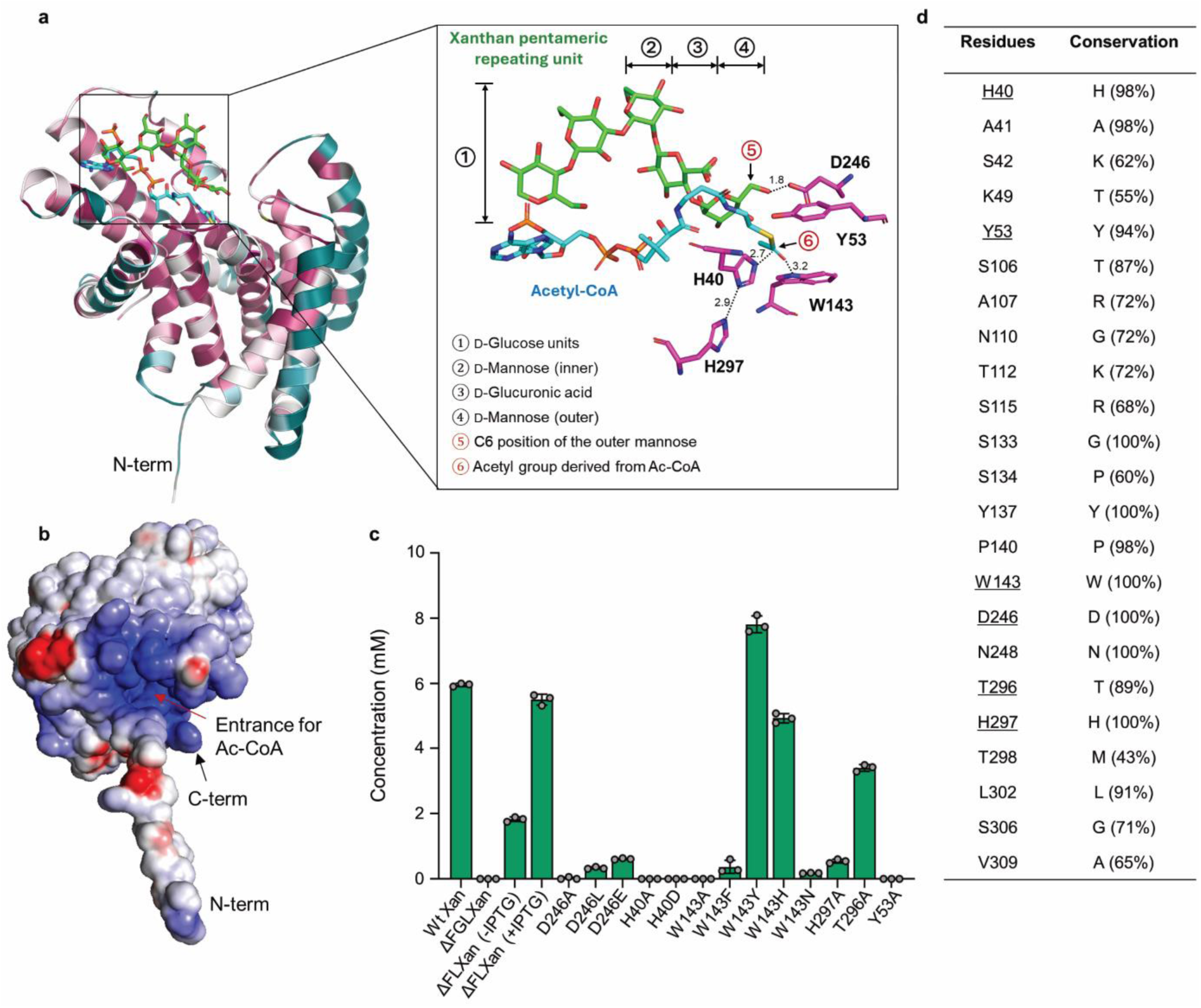
Structural and functional characterisation of GumG. **a,** ConSurf analysis of GumG in complex with docked acetyl-CoA (Ac-CoA, blue) and a xanthan pentameric repeat unit (green). Residues are coloured according to conservation scores (dark purple, highly conserved; blue, less conserved) based on a multiple sequence alignment of 49 GumG homologues. Catalytically important residues are highlighted, and inter-residue and residue–ligand distances were measured in PyMOL. **b,** Electrostatic surface potential of GumG calculated using the Adaptive Poisson–Boltzmann Solver (APBS). Surface charges are shown in blue (positive), red (negative) and white (neutral). A prominent positively charged patch is observed on the periplasmic surface and extending into the internal channel. **c,** Quantification of acetate in xanthan and xanthan variants produced by wild-type (WT) and engineered strains. Acetate content was determined by HPLC using a Rezex™ ROA H^+^ column following acid hydrolysis. Functional analysis of GumG was performed by plasmid-based complementation of the *Xcc ΔgumFGL* strain with an IPTG-inducible *pSRKGm* vector encoding GumG. Single amino acid substitutions were introduced by site-directed mutagenesis of *pSRKGm-GumG* and expressed in *Xcc ΔgumFGL*. **d,** Conservation analysis of residues surrounding the Ac-CoA binding pocket on the periplasmic face of GumG, based on a multiple sequence alignment of 49 homologues. Residues implicated in the proposed catalytic mechanism are underlined.

Surface electrostatic potential analysis using APBS revealed a prominent positively charged patch spanning the surface and internal tunnel of GumG, consistent with a potential Ac-CoA binding site mediated by interactions with the negatively charged phosphate groups of Ac-CoA (**Fig. 4b**). Complementary AutoSite analysis identified multiple putative binding pockets along the internal tunnel and adjacent solvent-exposed surface regions of GumG. Docking simulations with AutoDock Vina support a stepwise binding mode in which Ac-CoA binds GumG before the pentasaccharide substrate. Given that the pentasaccharide backbone is expected to be linked to undecaprenyl phosphate (Und-P), a lipid carrier embedded in the cell membrane, acetylation is likely to occur exclusively at the protein surface (**Fig. 1a**). Accordingly, predicted binding sites proximal to the TMH were excluded from further analysis.

Integrating electrostatic, conservation and docking analyses, two distinct binding sites were identified on GumG at the cytoplasmic and periplasmic sides. The cytoplasmic site lacks conserved catalytic residues and is therefore unlikely to support acetyl transfer, whereas the periplasmic site exhibits features consistent with a catalytically competent configuration and thus represents the most plausible locus for acetylation (**Fig. 4a**). These observations support a model in which Ac-CoA initially associates with the cytoplasmic side of GumG before translocating to the periplasmic functional site.

#### 3.3.1. Mechanistic insights into GumG catalysis based on mutagenesis

Based on the structural model of GumG with Ac-CoA and the pentameric xanthan unit bound at the periplasmic surface, conservation analysis of the 23 residues lining the Ac-CoA binding pocket revealed that His40, located within TMH1, is highly conserved (98%) across homologues and has been implicated in acetylation catalysis [9, 28, 29]. In addition, several proximal residues, including His297, Trp143, and Asp246, were fully conserved among the analysed ACTases, whereas Thr296 was conserved in approximately 89% of sequences, with the remaining positions occupied by Ala (6.5%) or Gly (4.3%) (**Fig. 4d**).

The functional contributions of these conserved residues were probed by alanine substitution. Replacement of His40, Asp246, Trp143, and His297 resulted in a marked reduction in xanthan acetylation, indicating their essential roles in GumG enzymatic activity and acetylation catalysis (**Fig. 4c**). The spatial arrangement of these key residues within the active site is consistent with a proposed catalytic mechanism for acetylation (**Fig. 5**). His40 and His297 are geometrically aligned, forming a near-linear hydrogen bond (∼180°) with a short donor–acceptor distance of 2.9 Å, consistent with proton transfer from His40 to His297 (**Fig. 5a**). The resulting deprotonated His40 is thus poised to act as a nucleophile, attacking the carbonyl carbon of Ac-CoA at a distance of 2.7 Å (**Fig. 5b**). The essential role of His297 is further supported by its high sequence conservation and the marked reduction of enzymatic activity to 9% following alanine substitution (**Fig. 4c,d**). Substitution of His40 with aspartic acid (H40D), a residue of comparable size but lacking nucleophilic character, completely abolished xanthan acetylation, confirming its essential catalytic role (**Fig. 4c**). The resulting tetrahedral intermediate is further stabilized by a hydrogen bond between the indole NH of Trp143 and the alkoxide oxgen (**Fig. 5b**). Mutagenesis of Trp143 revealed distinct functional contributions of aromatic side chains: substitutions with tyrosine (W143Y) or histidine (W143H) retained substantial acetylation activity (126% and 79%, respectively), whereas replacement with asparagine (W143N) drastically diminished activity to 3% (**Fig. 4c**). These observations underscore the importance of π-electron-rich aromatic residues in proton transfer and stabilization of reaction intermediates during catalysis. Finally, collapse of the tetrahedral intermediate, stabilised by Trp143, releases CoASH and forms an acetyl–His40 adduct (**Fig. 5c**).

**Fig. 5.**
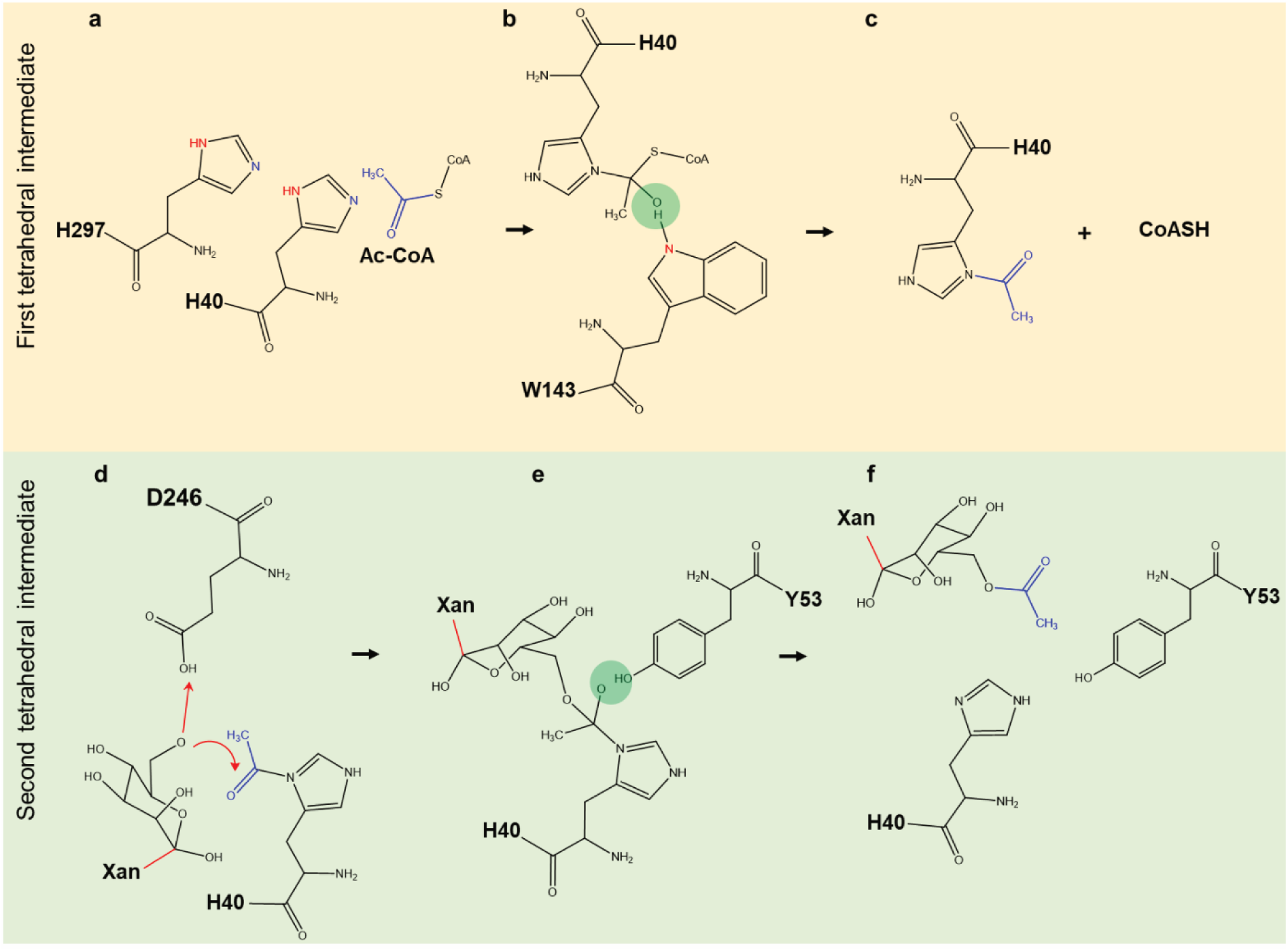
Proposed mechanism of GumG-mediated *O*-acetylation of a xanthan pentameric saccharide. Acetyl-CoA (Ac-CoA) is proposed to translocate from the cytoplasmic side to the periplasmic face of GumG. **a**, The alignment of His40 and His297 supports activation of His40 as a nucleophile, enabling nucleophilic attack on the thioester bond of Ac-CoA to form a tetrahedral intermediate. **b,** This intermediate is stabilised by hydrogen bonding with Trp143. and subsequently collapses to release CoASH and generate an acetylated His40 (Ac–His40) intermediate. **c,** Concurrently, a glycan strand of the lipid-anchored xanthan pentameric unit binds to the active site. Asp246, positioned on the protein surface, acts as a general base by abstracting a proton from the hydroxyl group of the outer mannose (Man) residue, predominantly at the C6 position. **d-e,** The resulting deprotonated hydroxyl group then attacks the carbonyl carbon of the Ac–His40 intermediate, forming a second tetrahedral intermediate that is potentially stabilised by hydrogen bonding with Tyr53. **f,** Collapse of this intermediate yields the *O*-acetylated pentameric saccharide and regenerates the free enzyme for subsequent catalytic cycles.

Concurrently, a glycan strand of the xanthan pentameric unit, anchored to the cell membrane via a lipid carrier, binds to the active site of GumG. Asp246, positioned on the protein surface, acts as a general base by abstracting a proton from the hydroxyl group of the outer Man residue in the xanthan repeating unit, primarily at the C6 position, with a proton transfer distance of 1.8 Å (**Fig. 5d**). This mechanistic role is supported by mutagenesis experiments: substitution of Asp246 with alanine completely abolished acetylation activity (**Fig. 4c**). Replacement with leucine (D246L), which introduces a nonpolar alkyl chain in place of the carboxylate, markedly impaired proton abstraction, reducing acetylation to 5% (**Fig. 4c**). Similarly, substitution with glutamic acid, which preserves the carboxylate but introduces a longer side chain and altered spatial orientation, decreased acetylation to 10% (**Fig. 4c**). Together, these results underscore the critical importance of side-chain length and spatial orientation for substrate positioning and catalytic efficiency. The resulting deprotonated hydroxyl group of the terminal Man nucleophilically attacks the carbonyl carbon of the acetyl-His40 intermediate, forming a second tetrahedral intermediate (**Fig. 5e**). This intermediate is stabilized by hydrogen bonding from Tyr53, which facilitates proton transfer, ultimately leading to release of the *O*-acetylated pentasaccharide and regeneration of the free enzyme (**Fig. 5f**). Consistent with this mechanism, substitution of Tyr53 with alanine abolished enzymatic activity (**Fig. 4c**).

#### 3.3.2. Rational engineering of GumG for expanded substrate selectivity

To improve GumG substrate selectivity through rational design, the cytoplasmic Ac-CoA binding site was selected as the engineering target. This site is proposed to function as an intracellular Ac-CoA binding site and entry channel, guiding CoA thioesters toward the periplasmic catalytic region (**Fig. 6a**). Within this site, 19 residues lining the Ac-CoA pocket were identified and subjected to systematic conservation analysis (**Fig. 6d**). To assess the functional contribution of each residue, single alanine substitutions were introduced at the corresponding positions.

**Fig. 6.**
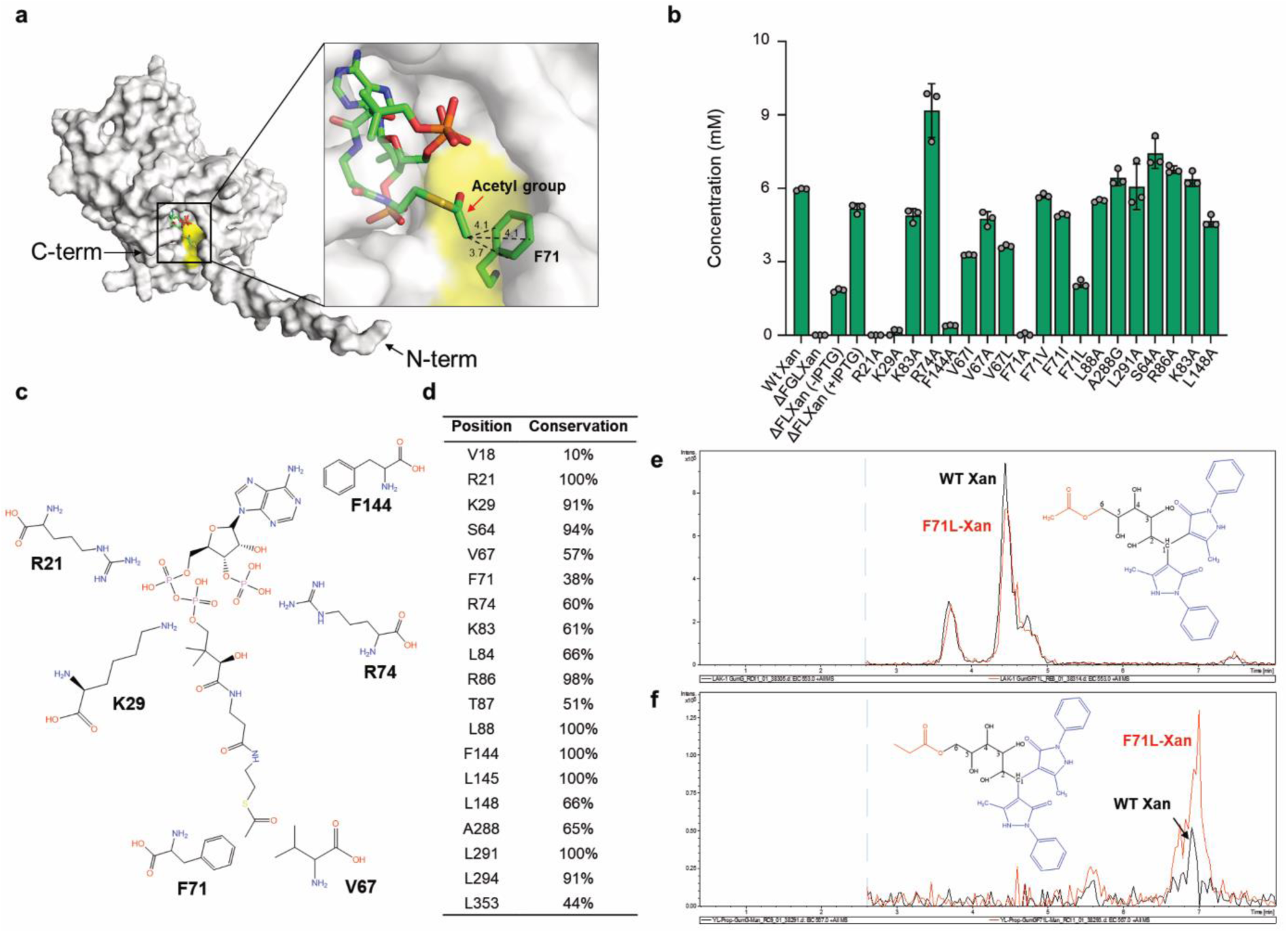
Structural and functional characterisation of GumG. **a,** Potential acetyl-CoA (Ac-CoA) binding pocket on the cytoplasmic face of GumG. Amino acid residues lining the acetyl group binding site are highlighted in yellow. **b,** Acetate levels in xanthan variants produced by the wild-type (WT) and engineered strains were quantified by HPLC using a Rezex™ ROA H⁺ column. The function of GumG was investigated by plasmid-based complementation of the *ΔgumFGL* mutant using the IPTG-inducible *pSRKGm* vector harboring the GumG gene. Single amino acid substitutions were generated by site-directed mutagenesis of *pSRKGm-GumG* and subsequently expressed in *Xcc ΔgumFGL*. For hydrolysis, polymer samples (10 g L⁻¹) were treated with 250 mM H₂SO₄ at 90 °C for 16 h. **c,** Key residues surrounding the Ac-CoA binding pocket. **d,** Conservation analysis of residues surrounding the Ac-CoA binding pocket, determined by multiple sequence alignment of GumG with 49 homologs. **e-f,** Activity assays of WT GumG (black) and the F71L GumG variant (red) using D-mannose (D-Man) as the acceptor and either **e,** Ac-CoA or **f,** propionyl-CoA (PA-CoA) as the donor. The structures of the *O*-6 acetylated and propionylated products are shown as representative examples, however, due to the catalytic promiscuity, modifications may also occur at additional positions. Extracted ion chromatograms (EIC, m/z 553.0 and m/z 567.0) represent acetylated and propionylated Man derivatives from WT GumG and the F71L GumG variant, respectively.

Conservation analysis revealed that Arg21 is fully conserved (100%), whereas Lys29 is conserved in 91.3% of sequences, with arginine present at this position in the remaining 8.7% of ACTases, indicating strong conservation across the enzyme family (**Fig. 6d**). Alanine substitution of these residues resulted in a marked reduction in polymer acetylation, decreasing to 5% in R21A-Xan and 8% in K29A-Xan, respectively (**Fig. 6b**). Structural modelling suggests that these positively charged residues stabilize the phosphopantetheine arm of Ac-CoA through ionic interactions (**Fig. 6c**). By contrast, although Arg74 was hypothesised to stabilize Ac-CoA through interactions with the phosphate group of the adenosine moiety, substitution of Arg74 with alanine did not reduce xanthan acetylation. This observation suggests that stabilization of the phosphopantetheine arm, rather than interactions with the adenosine phosphate, is the dominant determinant of Ac-CoA binding (**Fig. 6b, c**). In addition, Phe144, which exhibits lower sequence conservation, was found to be critical for substrate binding. Substitution of Phe144 with alanine reduced xanthan acetylation to 1% of WT GumG activity (**Fig. 6b, c**). Given its proximity to the adenine moiety of Ac-CoA, this effect is consistent with a π-stacking interaction between the aromatic side chain of Phe144 and the adenine ring that contributes to substrate stabilization.

Based on the structural model, Val67 and Phe71 were hypothesised to form a hydrophobic pocket accommodating the acetyl group (**Fig. 6a**). To preserve the hydrophobic character of the binding pocket while reshaping its geometry, Val67 and Phe71 were substituted exclusively with hydrophobic residues. Notably, sequence alignment revealed that both residues are poorly conserved among the evaluated ACTases (**Fig. 6d**). Substitution of Val67 with alanine, isoleucine or leucine reduced acetylation to 92%, 63%, and 70%, respectively, indicating that side-chain length at this position is an important determinant of GumG activity *in vivo* (**Fig. 6b**). By contrast, mutagenesis of Phe71 revealed a more complex structure-activity relationship. Replacement with alanine, valine, isoleucine or leucine resulted in acetylation levels of 1% (F71A), 110% (F71V), 95% (F71I), and 40% (F71L) relative to WT GumG (**Fig. 6b**). Together, these results indicate that both side-chain length and spatial orientation at this position critically influence GumG catalytic efficiency.

To further characterise the functional consequences of these mutations, *in vitro* assays were performed using a panel of GumG variants (K29A, R21A, V67A, V67I, V67L, F71A, F71V, F71I, and F71L) that exhibited pronounced effects on xanthan acetylation. All variants were heterologously expressed in *E. coli* C43 and evaluated for transferase activity in a one-pot reaction system coupling the ACTase assay with *in situ* generation of CoA thioesters, including PA-CoA, CA-CoA, AA-CoA, MAA-CoA. D-Man was selected as the acceptor substrate, as it displayed higher acetylation efficiency than the pentasaccharide acceptor *in vitro*. WT GumG efficiently transferred acetyl groups to D-Man, while exhibiting minimal activity toward PA-CoA (**Fig. 6e**). Notably, the F71L substitution altered substrate specificity, increasing activity toward PA-CoA while modestly reducing activity toward the native substrate Ac-CoA, as inferred from relative signal intensities, although accurate quantification was limited by the lack of appropriate standards (**Fig. 6e, f**). By contrast, the remaining variants lost propionyl-transfer activity but retained partial activity toward Ac-CoA. These results indicate that Phe71 plays a key role in shaping the initial Ac-CoA binding pocket, thereby influencing substrate selectivity and regulating access of Ac-CoA and related CoA thioesters to the catalytic site. Consistent with these observations, LC-MS/MS analysis failed to detect modified sugar products in reactions employing CA-CoA, AA-CoA or MAA-CoA generated via the one-pot enzymatic system.

## 4. Discussion

Microbial exopolysaccharides, such as xanthan, represent sustainable and biodegradable alternatives to petroleum-based polymers. Functionalization—particularly at hydroxyl groups—enables the generation of materials with tailored, application-specific properties. However, achieving such modifications through biocatalysis requires a deeper understanding of ACTases, such as GumG, and the rational engineering of these enzymes to enable efficient and versatile acyl transfer onto polysaccharides.

GumG, an ACTase that modifies the outer Man residues of xanthan side chains, poses substantial challenges for biochemical and structural characterisation owing to its membrane-associated nature, low solubility and limited stability *in vitro*. As a result, structural and mechanistic information on GumG and other members of the AT-3 family remains scarce. Here, an integrated analytical strategy combining site-specific enzymatic depolymerisation of xanthan with LC-MS/MS analysis was developed to characterise structural variants of xanthan produced by WT and genetically engineered *Xcc* strains. This approach enabled high-resolution mapping of site-specific modifications in complex microbial polysaccharides. To this end, RuGH5a, a truncated xanthanase from *Ruminococcaceae*, was employed to selectively hydrolyse xanthan at the reducing-end Glc residue of the backbone, irrespective of acetylation or pyruvylation status [19]. The resulting pentameric fragments were subsequently derivatised with PMP, which reacts with the open-ring form of the reducing sugar to yield stable bis-PMP adducts [24]. This derivatisation constrains MS/MS fragmentation to a single, well-defined pathway initiated from the reducing-end Glc, thereby enabling unambiguous structural elucidation of xanthan pentamers [24]. In parallel, a xanthan lyase from *Paenibacillus polymyxa* was used to selectively remove the outer Man residues of xanthan, independent of their acetylation or pyruvylation status, allowing targeted monitoring of modifications on these terminal Man residues [25].

Analysis of the outer Man residues in XanΔFL revealed four distinct peaks corresponding to *O*-2, *O*-3, *O*-4, and *O*-6 acetylation, with the *O*-1 and *O*-5 positions occupied by glycosidic linkages. Based on relative peak intensity, the dominant chromatographic peak was tentatively assigned to *O*-6 acetylation, using 6-*O*-acetyl-D-Glc as a surrogate for the unavailable 6-*O*-acetyl-D-Man standard; however, precise quantification of acetyl positional isomers was not possible in the absence of authentic acetylated mannose standards. Traditional analytical methods, such as NMR, offer limited sensitivity for resolving complex or low-abundance linkages and may therefore have overlooked acetylation at alternative positions [30]. These findings indicate that GumG exhibits spatially non-specific acetylation activity, challenging previous reports that xanthan acetylation is restricted to the *O*-6 position [31, 32]. In plant polysaccharides such as xylan, pectin and mannan, acetylation predominantly occurs at the *O*-2 and *O*-3 positions, with site variability influenced by hydroxyl accessibility and ACTase catalytic mechanisms [33–35]. Acetylation position has pronounced effects on polymer properties, for example, *O*-2/*O*-3 modification of β-mannan alters solubility and chain flexibility [36]. Notably, although multiple acetylation positions in xanthan were observed in earlier work by Ostrowski *et al.*[19], this feature was not explicitly discussed. In contrast, Stankowski et al. demonstrated the presence of a doubly acetylated outer mannose in xanthan [37].

GumG’s broad substrate specificity was confirmed *in vitro*, with acetylation observed for monosaccharides (D-Man, D-Glc, D-Gal), a disaccharide (Man_2_) and pentasaccharides (PenΔFGL, PenΔGL), whereas L-Man was not accepted, consistent with its rare biological occurrence [38, 39]. This promiscuity highlights the potential of GumG for the biocatalytic production of sugar esters, a class of biosurfactants, under aqueous conditions, thereby circumventing the solubility constraints associated with conventional chemical approaches [40]. Notably, *in vitro* acetylation of pentasaccharides preferentially targeted the non-reducing-end Glc residue, in contrast to the outer mannose acetylation pattern observed in native xanthan. This discrepancy likely reflects the constrained orientation of the lipid-linked pentasaccharide substrate *in vivo*, imposed by membrane anchoring through the Und-P carrier [41]. Consistent with this interpretation, Ac-CoA-dependent acetylation has been demonstrated on lipid-linked xanthan intermediates [42] and on semisynthetic peptidoglycan oligomers by the *Staphylococcus aureus O*-ACTase OatA [9], although positional specificity was not resolved in either case. Together, these observations underscore the challenges inherent in defining site-specific acetylation within complex polysaccharides and emphasise the need for integrated structural, biochemical and mechanistic analyses of ACTases such as GumG.

GumG also exhibited pronounced catalytic promiscuity, acting on diverse acceptors and two acyl-CoA donors, consistent with other AT3-domain enzymes capable of transferring alternative acyl groups, including succinate, isovalerate and propionate [16]. Comparable donor flexibility has been reported for the ACTase Wssi, which acetylates cellulose oligomers using both chemically activated acetyl donors and Ac-CoA [43]. By contrast, GumG displayed a more restricted donor spectrum, preferentially transferring short-chain CoA thioesters (≤C3), with substantially lower activity toward PA-CoA and AA-CoA relative to its native donor Ac-CoA. Direct quantitative comparison between PA-CoA and AA-CoA was further complicated by the intrinsic instability of AA-CoA, owing to the rapid degradation of its reactive vinyl group, which could not be mitigated by conventional stabilisers [44]. In this context, MAA-CoA represents a promising alternative acyl donor for the generation of UV-crosslinkable polymers [45]. However, its increased steric bulk is likely to necessitate targeted enzyme engineering to achieve efficient acyl transfer.

Although most models propose that xanthan acetylation occurs on the cytoplasmic side before the pentasaccharide is flipped for polymerisation, direct experimental support for this view remains limited [46]. Molecular dynamics simulations by Newman *et al.* suggest that Ac-CoA can span the membrane, positioning the acetyl group in proximity to extracytoplasmic catalytic residues [12]. This model is consistent with observations in other ACTases, including the extracytoplasmic SGNH domain of OatA, a peptidoglycan *O*-ACTase [9, 47].

Structural and functional analyses of GumG provide mechanistic insight into its catalytic activity, particularly with respect to Ac-CoA binding and substrate acetylation. Two conserved residues, Arg21 and Lys29, are likely to stabilise Ac-CoA through cationic interactions, analogous to arginine-mediated coordination of CoA phosphates in *Staphylococcus aureus* OatA (SaOatA). These residues correspond to the conserved Arg/Lys-X_10_-His motif (Lys29-His40 in GumG), supporting a central role in acetyl transfer catalysis [9, 16]. In addition, a π-stacking interaction between Phe144 and the adenine ring of CoA, also observed in SaOatA and *Nm*OatWY [9, 48], together with a hydrophobic pocket formed by Val67 and Phe71, contributes to ligand accommodation within the Ac-CoA binding site [49]. The latter feature is reminiscent of acyl-binding architectures observed in AMP-forming Ac-CoA synthetases. Consistent with these structural predictions, mutagenesis of these residues modulated xanthan acetylation, with the F71L substitution enhancing PA-CoA utilisation while modestly reducing activity toward the native donor Ac-CoA. Comparable structure-guided engineering strategies have been reported for GNAT family enzymes and chloramphenicol ACTases, in which single amino-acid substitutions alter acyl-donor specificity and expand the range of utilisable acyl-CoA substrates [50, 51]. On the periplasmic face of GumG, a putative catalytic triad includes His40, a conserved residue within TMH1 that is frequently implicated in catalysis among AT3-domain enzymes [9, 16]. In *Streptococcus thermophilus* OatA, histidine functions as a general base to activate a tyrosine nucleophile [9], whereas in lysosomal Ac-CoA:α-glucosaminide *N*-ACTase it acts directly as the catalytic nucleophile [28, 29]. These mechanistic precedents highlight the functional versatility of conserved histidine residues in acyltransferase catalysis.

While ACTases typically employ an acid-base-nucleophile catalytic triad, some enzymes, such as carnitine ACTase, proceed through two tetrahedral intermediates, with a deprotonated substrate hydroxyl acting as a second nucleophile [9, 15]. In GumG, the spatial arrangement of conserved residues is consistent with a related dual-triad mechanism, in which Asp246 on the protein surface may function as a general base to promote nucleophilic attack on the outer Man hydroxyl group of xanthan. Although histidine-mediated proton transfer involving a tyrosine residue has been proposed for other ACTases [9], the distances between His40, Tyr53 and the carbonyl carbon in the GumG structural model are incompatible with such a pathway, arguing against a tyrosine-activated nucleophilic mechanism in this case.

Together, these findings provide a mechanistic basis for understanding how acyl-donor selectivity is encoded in membrane-associated ACTases and offer design principles for tuning microbial polysaccharide modification.

## Acknowledgments

This work was funded through the Perfecoat project titled “High performance Bio-based Functional Coatings for Wood and Decorative Applications” (Grant 101022370).

## Author contributions

Y.L., B.R., T.B., and M.D. analyzed and interpreted the data. O.M. and Y.L performed protein modeling and molecular docking. Y.L. designed the experiments and acquired the data. L.K. contributed to part of the *in vitro* experiment. Y.L. drafted the manuscript, which was revised by V.S., T.B. and B.R.

