## Supplementary Data for "Unraveling *Xanthomonas* acetyltransferase GumG: decoding catalytic promiscuity, understanding the mechanism, and enhancing enzymatic versatility"

#### Supplementary Fig.1

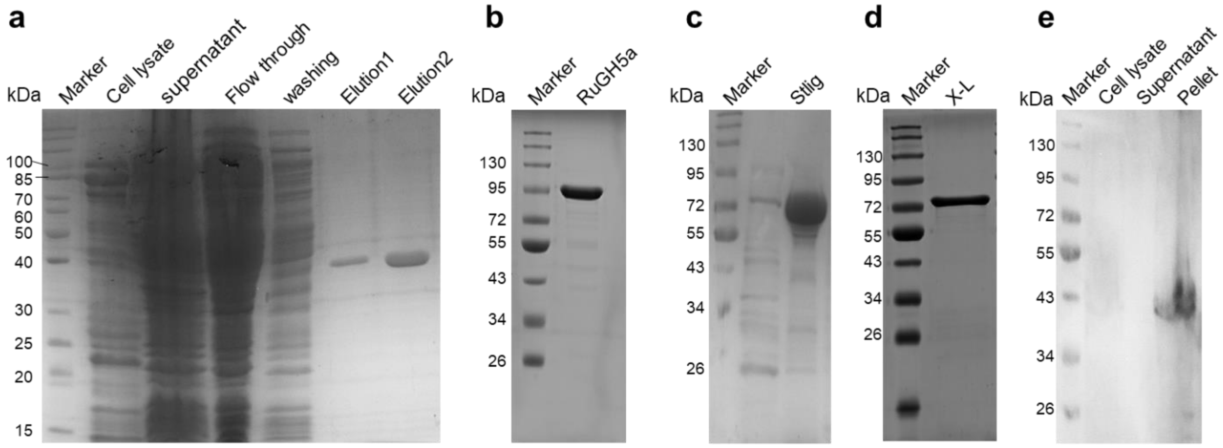


**Supplementary Fig. 1 | Protein expression and purification. a,** Expression and purification of recombinant GumG in *E. coli* C43 by immobilised metal affinity chromatography (IMAC) on a Ni–TED column. Lane 1, prestained protein standards; lane 2, whole-cell lysate after ultrasonication; lane 3, clarified supernatant after centrifugation; lane 4, flow-through fraction; lane 5, wash fraction; lanes 6–7, elution fractions. **b-c,** Expression and purification of **b,** xanthanase RuGH5a (95.3 kDa) and **c,** Stlig (75.0 kDa) in *E. coli* BL21(DE3). **d,** SDS–PAGE analysis of commercial xanthan lyase (X-L, 81.6 kDa) from *Bacillus sp*. (Megazyme). **e,** Western blot analysis of GumG expression (40.0 kDa) and subcellular localisation in *Xcc ΔgumFGL* using an anti-His antibody. Lane 1, Whole-cell lysate; lane 2, soluble supernatant; lane 3, membrane-containing pellet fractions. Molecular weight markers (kDa) are indicated on the left. For heterologous expression in *E. coli*, proteins were expressed from *pET28*-based constructs. For expression in *Xcc ΔgumFGL*, GumG was cloned into the *pSRKGm* vector. All constructs carried a C-terminal $His_6$ tag for purification and detection.

**Supplementary Fig.2**

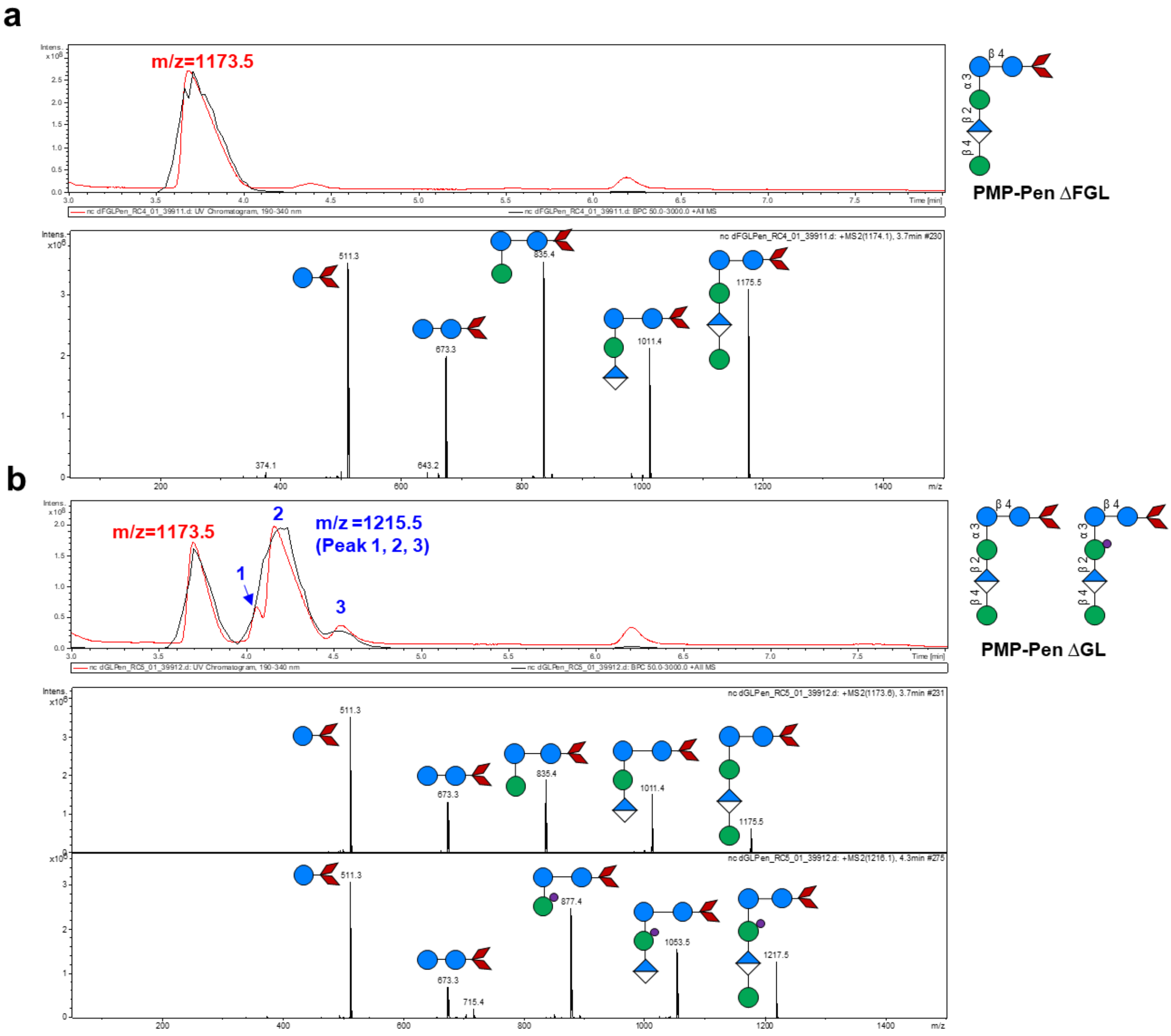


**Supplementary Fig. 2 | Extracted ion chromatograms and mass spectra of RuGH5a digestion products from XanΔFGL and XanΔGL. a,** UV chromatogram, base peak chromatogram (BPCs) and the corresponding MS/MS spectra of RuGH5a digestion products derived from XanΔFGL. The chromatographic peak at *m/z* 1173.5 corresponds to PMP-derivatised non-acetylated pentameric fragments (PenΔFGL). The MS/MS spectrum of the *m/z* 1173.5 ion shows a series of fragment ions at *m/z* 511.3, 673.3, 835.4, 1011.4 and 1175.5, corresponding to the PMP-derivatised monomer through pentamer, respectively. **b,** UV chromatogram, BPC and the corresponding MS/MS spectra of RuGH5a digestion products from XanΔGL. Co-elution of peaks at *m/z* 1173.5 and 1215.5 indicates the presence of non-acetylated and singly acetylated pentameric fragments. The *m/z* 1173.5 species exhibits an identical fragmentation pattern to PenΔFGL, confirming the presence of non-acetylated pentamers. The *m/z* 1215.5 species comprises three distinct chromatographic peaks (peaks 1–3), consistent with isomeric forms of singly acetylated pentamers exhibiting different hydrophobicities. MS/MS spectra from all three peaks display fragment ions at *m/z* 511.3 (monomer), 673.3 (dimer), 877.4 (acetylated trimer), 1053.5 (acetylated tetramer) and 1217.5 (acetylated pentamer). Pentameric xanthan oligosaccharides were generated by digestion of xanthan (5 g $L^{-1}$) with RuGH5a (1.05 µM) at 37 °C for 72 h, followed by isopropanol precipitation and purification by Bio-Gel P2 size-exclusion chromatography using an ÄKTA system with water as the eluent. Fractions containing pentameric products were identified by DNS assay and LC-MS/MS, pooled and concentrated by SpeedVac prior to analysis. **Symbol legend:** red rhombus, PMP; blue circle, glucose (Glc); green circle, mannose (Man); blue-and-white diamond, glucuronic acid (GlcA); purple circle, acetate (Ac).

#### Supplementary Fig.3

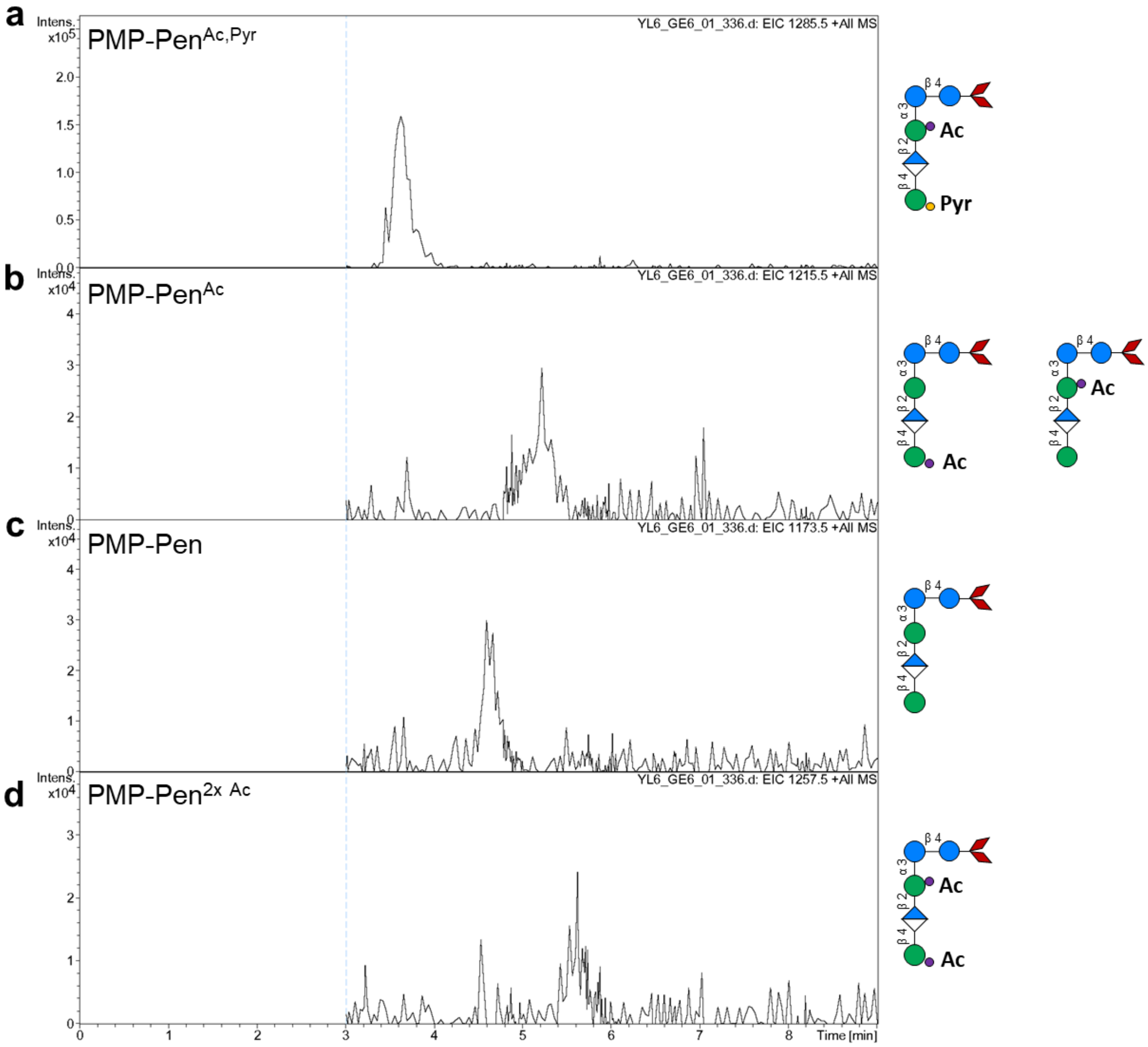


**Supplementary Fig. 3 | Extracted ion chromatograms of RuGH5a digestion products from wild-type xanthan.** Extracted ion chromatograms (EICs) of pentameric products generated by RuGH5a digestion of wild-type xanthan (WT Xan). **a–d,** Four major pentameric species were detected: **a,** *m/z* 1285, corresponding to a pentamer bearing an acetyl group on the internal mannose (Man) residue and a pyruvyl group on the terminal Man; **b,** *m/z* 1215.5, corresponding to a singly acetylated pentamer with the acetyl group located on either the internal or terminal Man; **c,** *m/z* 1173.5, corresponding to a non-modified pentamer; and **d,** *m/z* 1257.5, corresponding to a doubly acetylated pentamer bearing acetyl groups on both the internal and terminal Man residues. Pentameric xanthan oligosaccharides were generated by digestion of xanthan (5 g $L^{-1}$) with RuGH5a (1.05 µM) at 37 °C for 72 h, followed by isopropanol precipitation, SpeedVac drying and resuspension in water prior to analysis. **Symbol legend:** red rhombus, PMP; blue circle, glucose (Glc); green circle, mannose (Man); blue-and-white diamond, glucuronic acid (GlcA); purple circle, acetate (Ac); yellow circle, pyruvate (Pyr).

#### Supplementary Fig.4

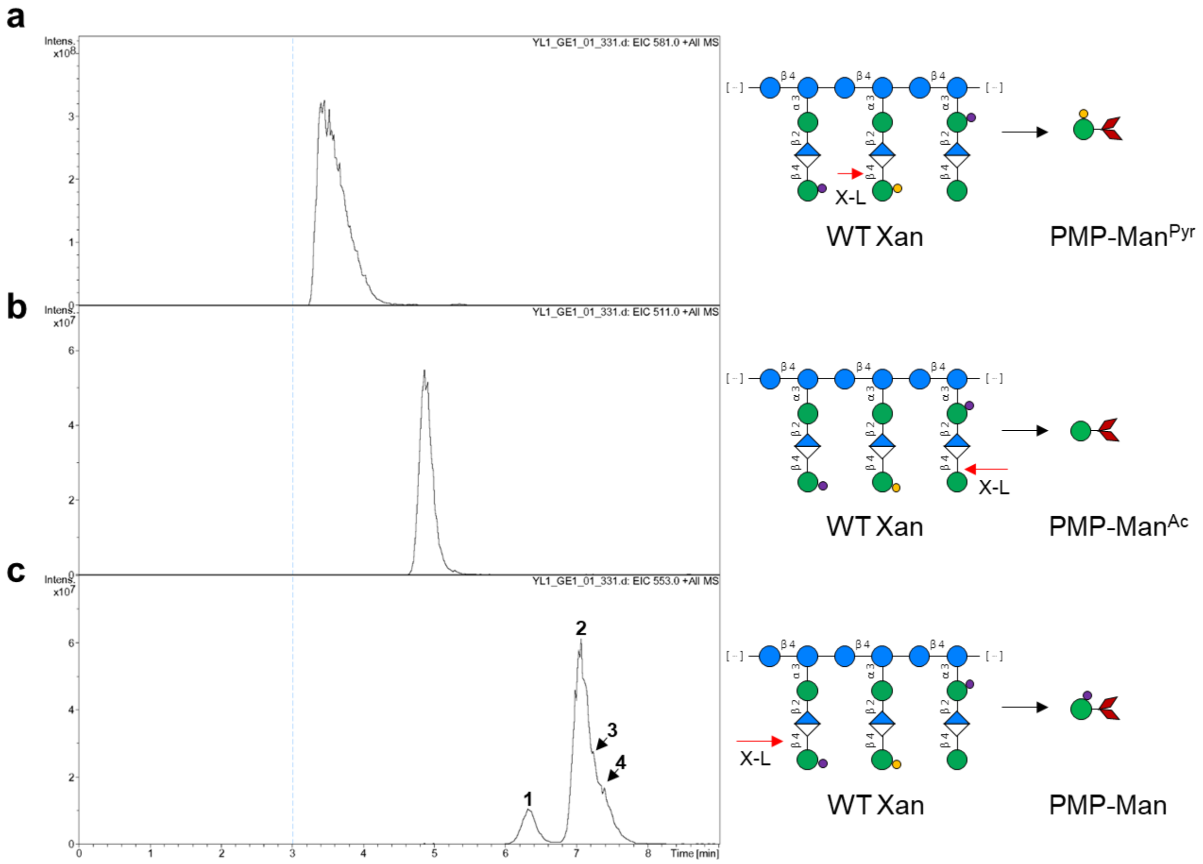


**Supplementary Fig. 4 | Extracted ion chromatograms of enzymatic digestion products from wild-type xanthan treated with xanthan lyase.** Extracted ion chromatograms (EICs) of monomeric products generated from the digestion of wild-type xanthan (WT Xan) by xanthan lyase (X-L). **a–c,** Three monomeric fragments corresponding to the outer mannose (Man) residue were detected: **a,** *m/z* 581, corresponding to a pyruvylated outer Man; **b,** *m/z* 511, corresponding to a non-acetylated outer Man; and **c,** *m/z* 553, corresponding to an acetylated outer Man. Outer Man residues were generated by digestion of WT xanthan (5 g $L^{-1}$) with X-L (101.54 nM) at 37 °C for 72 h, followed by isopropanol precipitation, SpeedVac drying and resuspension in water prior to analysis. **Symbol legend:** red rhombus, PMP; blue circle, glucose (Glc); green circle, mannose (Man); blue-and-white diamond, glucuronic acid (GlcA); purple circle, acetate (Ac); yellow circle, pyruvate (Pyr).

#### Supplementary Fig.5

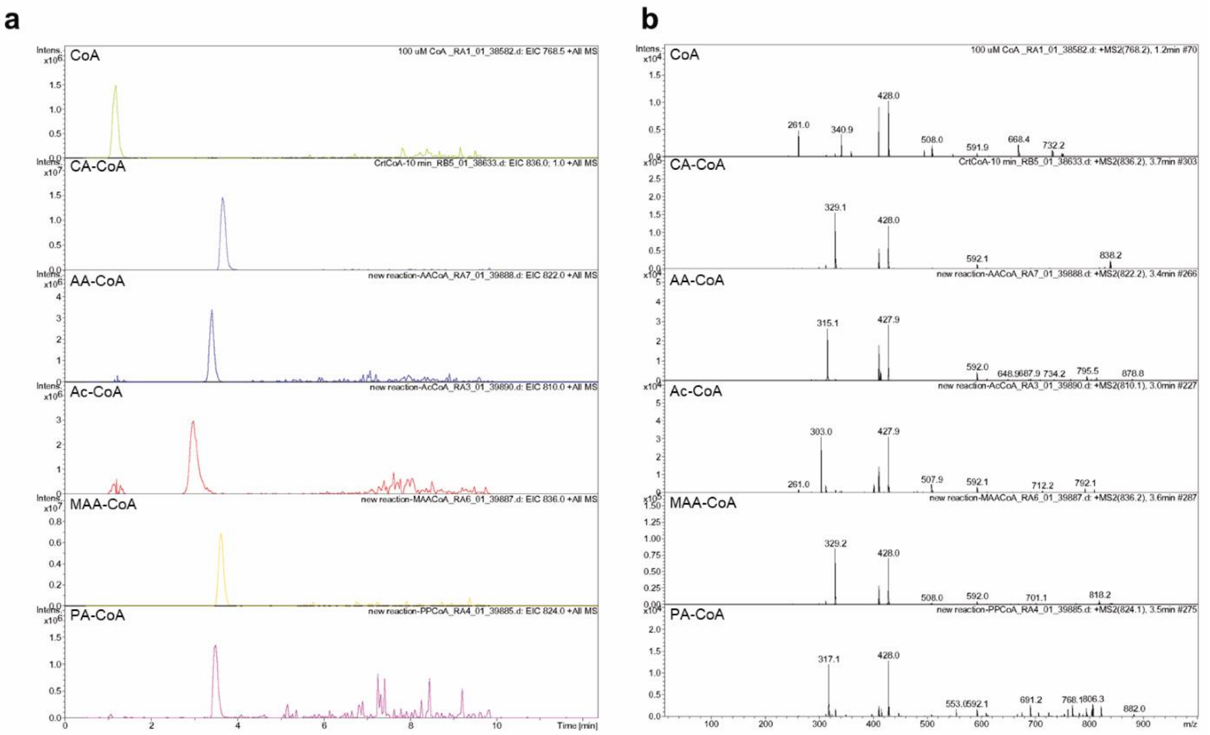


**Supplementary Fig. 5 | Extracted ion chromatograms and MS² spectra of CoA thioesters generated by Stlig.** **a,** Extracted ion chromatograms (EICs) of CoA thioesters produced by 3-hydroxypropionyl-coenzyme A synthetase (Stlig). Peaks at *m/z* 836.1, 768.5, 822.0, 810.0, 836.0 and 824.0 correspond to crotonyl-CoA (CA-CoA), coenzyme A (CoA), acryloyl-CoA (AA-CoA), acetyl-CoA (Ac-CoA), methacryloyl-CoA (MAA-CoA) and propionyl-CoA (PA-CoA), respectively. **b,** MS² fragmentation spectra of the corresponding CoA thioesters. Each species exhibited a characteristic neutral loss of 507 Da, yielding diagnostic fragment ions at *m/z* 329 for CA-CoA, *m/z* 261.5 for CoA, *m/z* 315 for AA-CoA, *m/z* 303 for Ac-CoA, *m/z* 329 for MAA-CoA and *m/z* 317 for PA-CoA. CoA thioesters were enzymatically synthesised in reaction mixtures containing 20 mM of the respective carboxylic acid (pH adjusted to 7.0 with NaOH), 0.25 mM CoA, 3 mM ATP, 10 mM $MgCl_2$ and 100 mM Tris–HCl buffer (pH 7.0), supplemented with 271.75 µM recombinant Stlig. Reactions were incubated at 37 °C for 10 min prior to LC-MS/MS analysis.

#### Supplementary Fig.6

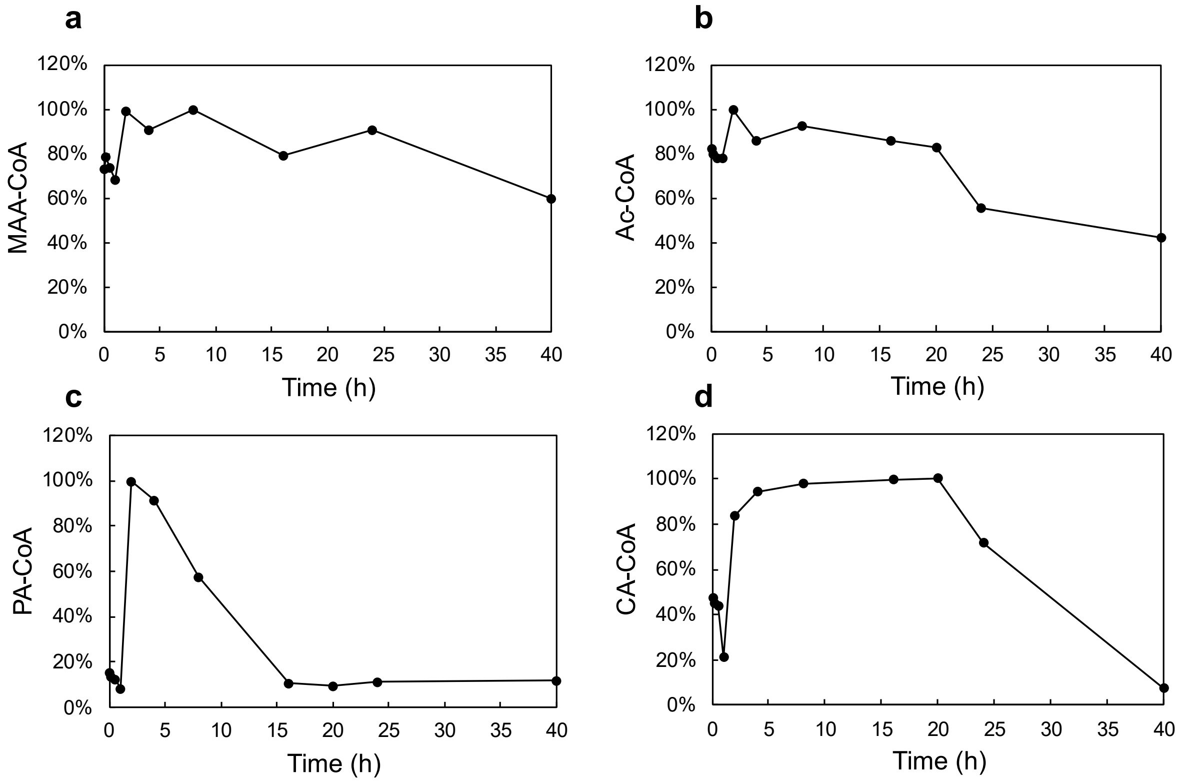


**Supplementary Fig. 6 | Time-course analysis of the stability of CoA thioesters.** Time-course analysis of methacryloyl-CoA (MAA-CoA), acetyl-CoA (Ac-CoA), propionyl-CoA (PA-CoA) and crotonyl-CoA (CA-CoA) generated by Stlig-catalysed reactions. **a-d,** Correspond to the compounds listed above. Relative abundances were calculated from peak areas of the corresponding CoA thioesters using Bruker Quant Data Analysis software. Each thioester was identified based on its characteristic *m/z* value and $MS^2$ fragmentation pattern. Monitoring of acryloyl-CoA (AA-CoA) over time was not feasible owing to its susceptibility to hydration at the carbon-carbon double bond, leading to rapid formation of hydroxypropionic acid within minutes. This assignment is supported by the characteristic ion at m/z 840 (derived from m/z 822 following addition of a water molecule, +18) and its corresponding fragment ion at m/z 315.

#### Supplementary Fig.7

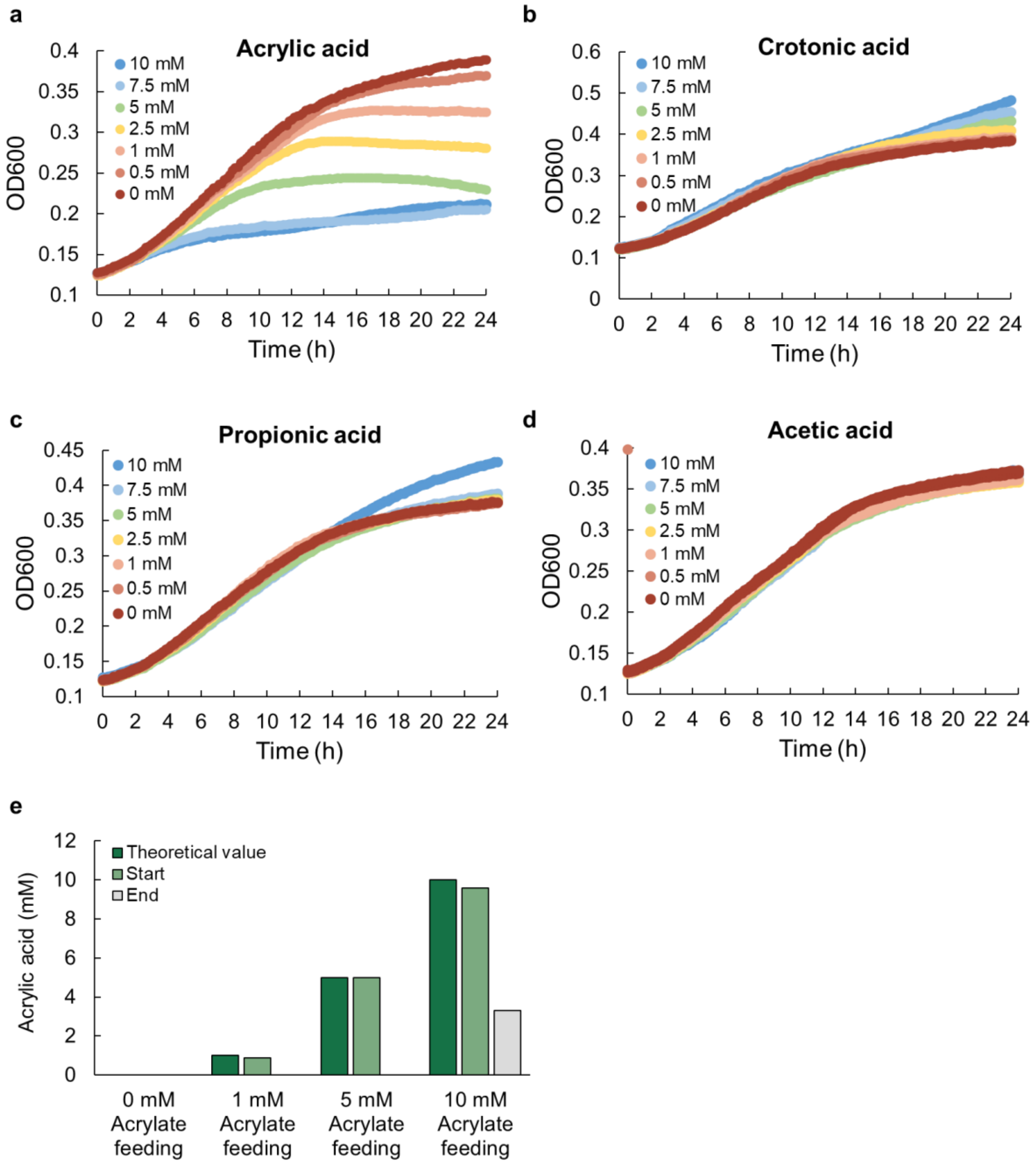


**Supplementary Fig. 7 | Growth profiles of *Xcc ΔgumL (pStlig)* in the presence of short-chain fatty acids and acrylic acid consumption.** Time-course growth profiles of *Xcc ΔgumL (pStlig)* cultured in LB medium supplemented with increasing concentrations of short-chain fatty acids: **a,** acrylic acid; **b,** crotonic acid; **c,** propionic acid; and **d,** acetic acid. Cultivations were initiated at an OD600 of 0.1 in 96-well plates containing 200 µl LB medium supplemented with 30 µg $mL^{-1}$ gentamicin and 1 mM IPTG. The indicated concentrations of the respective short-chain fatty acids were added, and growth was monitored in triplicate using a plate photometer. **e,** Assimilation of acrylic acid by cells at the beginning and end of cultivation under different feeding concentrations. As acrylic acid has been reported to interfere with the metabolic pathways of *E. coli*, consistent with our observations, its concentration in the culture was quantified to monitor its assimilation throughout the *in vivo* biotransformation process. Analysis was performed by HPLC using a Rezex$^{TM}$ ROA $H^+$ column. For this experiment, 30 mL cultures were inoculated at an OD600 of 0.1 and cultivated in 100 mL shaking flasks containing LB medium at 30 °C for 24 h with agitation at 300 rpm.

#### Supplementary Fig.8

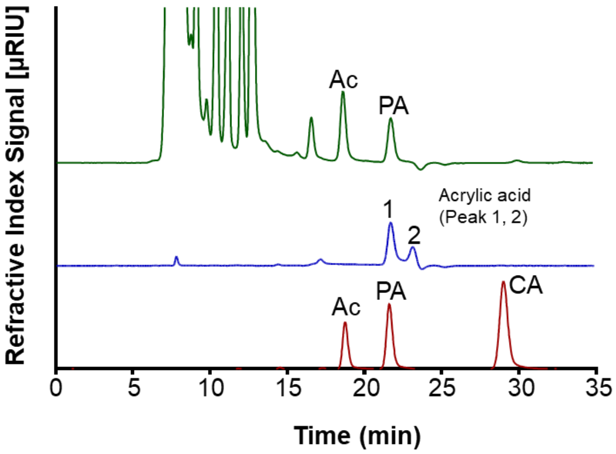


**Supplementary Fig. 8 | HPLC analysis of acrylic acid standards and xanthan hydrolysates.** HPLC analysis of acrylic acid standards and xanthan hydrolysates. The acrylic acid standard (blue trace) exhibited two peaks: a major peak co-eluting with propionic acid (peak 1) and a minor peak (peak 2) corresponding to intact acrylic acid. The red trace represents a standard mixture containing 5 mM each of acetate (Ac), propionate (PA) and crotonic acid (CA). Xanthan (10 g $L^{-1}$) spiked with 5 mM acrylic acid was subjected to acid hydrolysis in 250 mM $H_2SO_4$ at 90 °C for 16 h (green trace). The absence of acrylic acid (with the same retention time as peak 2) after hydrolysis indicates the loss of intact acrylic acid under these conditions, consistent with modification of the α,β-unsaturated C=C double bond during sample processing.

#### Supplementary Fig.9

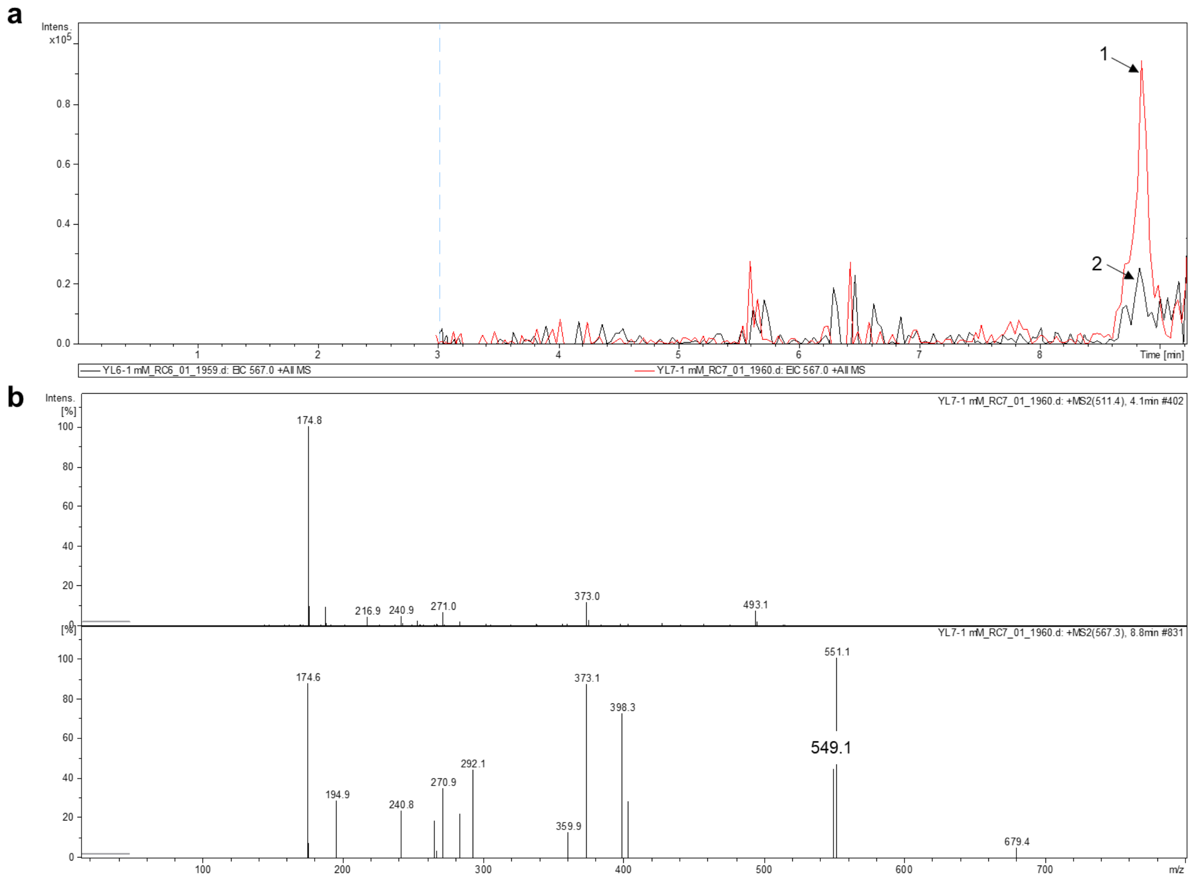


**Supplementary Fig. 9 | Analysis of outer mannose residues from xanthan produced by *Xcc ΔgumL (pStlig)* cultured with propionic acid. a,** Extracted ion chromatograms of outer mannose (Man) residues released from xanthan produced by *Xcc ΔgumL (pStlig)* (red trace) and by *Xcc ΔgumL* (black trace) cultured in the presence of 10 mM propionic acid. A prominent peak at *m/z* 567 corresponds to propionylated outer Man in polymer produced by *Xcc ΔgumL (pStlig)*. In contrast, xanthan from the control strain (*Xcc ΔgumL* lacking Stlig) exhibited only a weak signal at *m/z* 567. **b,** $MS^2$ spectra of outer Man residues. The propionylated Man species displayed a diagnostic fragment ion at *m/z* 549.1, corresponding to a +56 Da mass shift relative to non-acetylated Man (m/z 493.1), consistent with the propionylation of outer Man. Prior to analysis, xanthan samples were washed and dried three times with water to remove residual free acids. Outer Man residues were enzymatically released by digestion of xanthan (5 g $L^{-1}$) with xanthan lyase (X-L; 101.54 nM), followed by derivatisation with PMP.

### Supplementary Fig.10

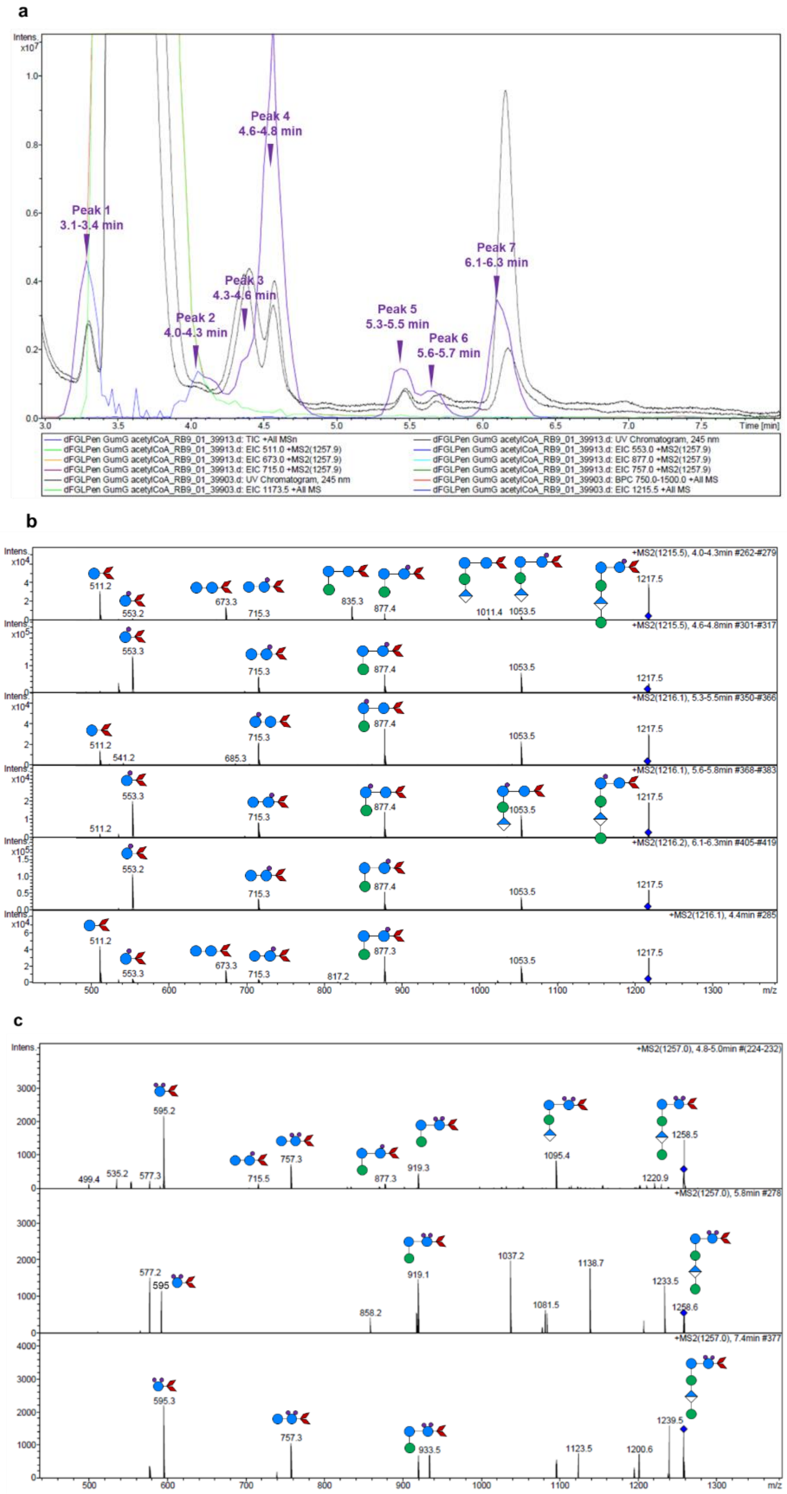


**Supplementary Fig. 10 | LC-MS/MS analysis of acetylated pentameric products generated from enzymatic reactions using PenΔFGL as the acceptor and acetyl-CoA as the donor. a,** UV chromatographic analysis and extracted ion chromatograms of singly acetylated PenΔFGL. Seven distinct UV-absorbing peaks at *m/z* 1215 were detected, consistent with the presence of multiple isomeric acetylated pentamers. **b,** MS/MS spectrum of singly acetylated PenΔFGL (*m/z* 1215), showing fragmentation patterns consistent with acetylation at different positions within the pentameric structure. **c,** MS/MS spectrum of doubly acetylated PenΔFGL (*m/z* 1257.0), confirming the presence of two acetyl groups and indicating distinct acetylation sites within the pentamer. One-pot enzymatic reactions were performed by incubating GumG (7.85 μM) and Stlig (3.63 μM) with CoA (250 μM), carboxylic acid (20 mM, pH adjusted to 7.0), ATP (3 mM) and PenΔFGL (13.67 mM) in 50 mM phosphate buffer (pH 7.0)

containing 200 mM NaCl and 10 mM $MgCl_2$ at 30 °C with shaking at 600 rpm for 48 h. Negative control reactions were performed by replacing GumG with buffer. The concentration of PenΔFGL was quantified using the DNS assay. **Symbol legend:** red rhombus, PMP; blue circle, glucose (Glc); green circle, mannose (Man); blue-and-white diamond, glucuronic acid (GlcA); purple circle, acetate (Ac).

#### Supplementary Fig.11

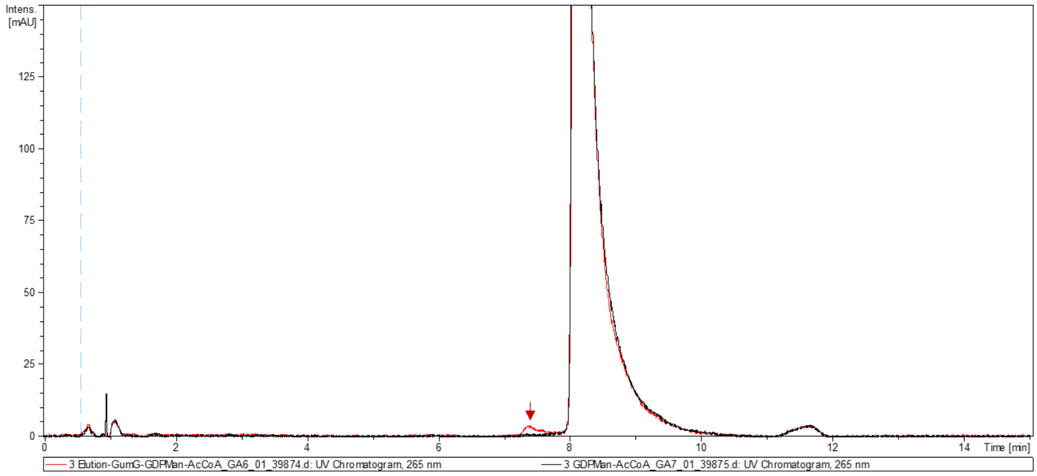


**Supplementary Fig. 11 | LC–MS/MS HILIC analysis of GumG-catalysed acetyl transfer to GDP-mannose.** LC–MS/MS HILIC analysis of products from GumG-catalysed transferase reactions using acetyl-CoA (Ac-CoA) as the donor and GDP-mannose (GDP-Man) as the acceptor. The red trace corresponds to reactions containing GumG, whereas the black trace represents the negative control in which GumG was replaced with buffer. A distinct peak at *m/z* 647 (arrow) was detected exclusively in reactions containing GumG and corresponds to acetylated GDP-Man. The peak at *m/z* 605, observed in both reaction and control samples, corresponds to non-acetylated GDP-Man. Reactions were performed by incubating Ac-CoA (50 mM) with GDP-Man (275 mM) in the presence of GumG (8.25 µM) at 30 °C for 48 h prior to LC–MS/MS analysis.

**Supplementary Method 1. Identifying the expression of GumG in *Xanthomonas campestris* Δ*gumFGL (Xcc* Δ*gumFGL) by* Western Blot**

To investigate the expression and subcellular localisation of plasmid-encoded full-length GumG bearing a C-terminal $His_6$ tag in *Xcc* Δ*gumFGL*, cells harbouring *pSRKGm-GumG* were induced with 1 mM IPTG at an $OD_{600}$ of 1.0 and cultured at 30 °C for 3 days. Cells from 1 mL cultures were harvested by centrifugation at 14,000 rpm for 1 min at 4 °C, washed twice with ice-cold 50 mM phosphate buffer (pH 7.0), and resuspended in 500 µL of the same buffer. Cell lysis was performed by ultrasonication on ice for 5 min (pulse duration 0.6 s; amplitude 60%). Membrane-containing fractions were subsequently collected by centrifugation at 14,000 rpm for 5 min at 4 °C, and the resulting pellets were resuspended in 500 µL phosphate buffer.

Total cell lysate, membrane-containing pellet and soluble supernatant fractions were separated by SDS-PAGE using 12% polyacrylamide gels and subsequently transferred onto polyvinylidene fluoride (PVDF) membranes. $His_6$-tagged proteins were detected using a mouse anti-Tetra-His primary antibody (Qiagen; 1:1,000 dilution) followed by a rabbit anti-mouse IgG (Fc)-conjugated alkaline phosphatase secondary antibody (Thermo Fisher; 1:1,000 dilution). Immunoreactive bands corresponding to GumG were visualised using BCIP/NBT substrate solution, prepared by diluting 66 µL NBT stock solution (Dissolve 1 g NBT dissolved in 20 mL of 70% DMF) and 33 µl BCIP stock solution (1 g BCIP dissolved in 20 mL of 100% DMF) in 10 mL Buffer A.

**Supplementary Method 2. Site-directed mutagenesis and plasmid construction**

Site-directed mutagenesis was performed using a pair of primers carrying the desired mutation at the 5′ end of the forward primer [1]. The entire plasmid was amplified using Phusion® High-Fidelity DNA polymerase. The resulting PCR product was treated with DpnI at 37 °C for 2 h to digest the methylated parental template, followed by enzyme inactivation at 80 °C for 10 min. Subsequently, 16 µL of the reaction mixture was incubated with 1 µL polynucleotide kinase (PNK), 1 µL T4 DNA ligase and 1 µL 10X T4 DNA ligase reaction buffer (all from New England Biolabs) at 37 °C for 1 h to enable simultaneous phosphorylation and ligation.

In addition, a modified mutagenesis strategy based on Gibson assembly was employed [2]. Two PCR fragments containing 20–25 bp homologous overlapping ends were generated independently, with the desired mutation introduced in the centre of one primer. These fragments were subsequently assembled using a Gibson assembly reaction according to the manufacturer's protocol.

For both approaches, up to 10 µL of the ligation or Gibson assembly reaction mixture was used to transform chemically competent *Escherichia coli* (*E. coli*) DH5α cells. Transformants were selected on LB agar plates supplemented with the appropriate antibiotics (30 µg $mL^{-1}$ gentamicin for plasmid *pSRKGm* and 100 µg $mL^{-1}$ kanamycin for plasmid *pET28*). Correct plasmid constructs were confirmed by Sanger sequencing (Eurofins Genomics) prior to transformation into expression hosts (*Xcc* Δ*gumFGL* or *E. coli* C43).

**Supplementary Method 3. Metabolite quenching and extraction of intracellular CoA and CoA thioesters from *Xcc***

A 50 mL culture of *Xcc ΔgumL* (*pStlig*) was grown in baffled shaking flasks at 30°C. At mid-exponential phase ($OD_{600}$ = 2.0–4.5), 10 mL of culture was harvested by centrifugation at 4,000 × g for 10 min at 4°C. Cell pellets were immediately placed on ice and resuspended in 2 mL of pre-cooled extraction solvent consisting of acetonitrile, methanol and water (40:40:20, v/v/v) at -20 °C. Metabolite quenching and extraction were performed for 1 h following the protocol described by Gansbiller et al. [3].

For each sample, 300 µL of the extract was dried using a SpeedVac concentrator. Dried residues were resuspended in 300 µL of water and filtered through a 0.22-µm PVDF syringe filter prior to liquid chromatography–tandem mass spectrometry (LC-MS/MS) analysis. For spiking experiments, 100 µM coenzyme A (CoA) or acetyl-CoA (Ac-CoA) was added either before or after solvent evaporation to assess compound stability and recovery.

**Supplementary Result 1. Structure analysis of xanthan and its variants reveals spatially non-specific acetylation by acetyltransferases**

To elucidate the catalytic specificities of the acetyltransferases (ACTases) GumG and GumF involved in xanthan biosynthesis, a series of xanthan variants—including wild-type xanthan (WT Xan), XanΔFL, XanΔGL, and XanΔFGL—were generated from WT *Xcc* and corresponding gene knockout strains (*Xcc*

*ΔgumFL*, *Xcc ΔgumGL, Xcc* Δ*gum*FGL), depolymerised and analysed. Structural analysis of XanΔFL provided direct evidence supporting the role of GumG in acetylation of the terminal (outer) Mannose (Man) residue, **as detailed in the manuscript**. In parallel, analysis of XanΔGL yielded insights into the function of GumF, which is proposed to acetylate internal Man residues. Analysis of WT Xan further suggests a synergistic interplay among GumG, GumF and GumL in modulating xanthan modification, with consequent effects on its structure and physicochemical properties.

Digestion of XanΔGL with the xanthanase RuGH5a yielded two major species at m/z 1173.5 and 1215.5, corresponding to non-acetylated and singly acetylated pentamers, respectively, consistent with partial mono-acetylation (+42 Da) catalysed by GumF. In contrast, digestion of XanΔFGL produced a single peak at m/z 1173.5, corresponding exclusively to non-acetylated pentamers. $MS^2$ analysis of the PenΔGL species at m/z 1215.5 revealed fragment ions at m/z 511.3, 675.3, 877.4, 1053.5 and 1215.5, corresponding to PMP-derivatised monomer, dimer, acetylated trimer, tetramer and pentamer fragments, respectively (**Fig. S2b**). By comparison, PenΔFGL displayed only non-acetylated fragment ions spanning monomer to pentamer, supporting the assignment of GumF as the enzyme responsible for acetylation of the internal Man residue (**Fig. S2a**). Furthermore, the presence of three distinct UV peaks within a single extracted ion chromatogram (EIC) at m/z 1215.5 indicates the existence of multiple positional isomers, suggesting that GumF acetylates the internal Man residue at three distinct hydroxyl positions, most likely at *O*-3, *O*-4 and *O*-6, as the remaining positions participate in glycosidic linkages (**Fig. S2b**).

In contrast, analysis of WT xanthan–derived pentamers revealed four prominent peaks at m/z 1173.5, 1215.5, 1257.5 and 1285.5 (**Fig. S3**). These peaks correspond to non-substituted pentamers (m/z 1173.5); singly acetylated pentamers (m/z 1215.5), in which the acetyl group is located on either the terminal or internal Man residue; doubly acetylated pentamers bearing one acetyl group on each Man (m/z 1257.5); and pentamers containing both an acetyl group (+42 Da) on the internal Man and a pyruvyl group (+70 Da) on the terminal Man (m/z 1285.5).

To support the structural assignments derived from pentamer analysis, outer Man residues from different xanthan variants were further released from the polymer backbone using xanthan lyase (X-L). As **described in the manuscript,** analysis of XanΔFL revealed both partially acetylated (m/z 553) and non-acetylated (m/z 511) outer Man residues **(Fig. 1d).** By contrast, only the non-acetylated Man PMP derivative (m/z 511) was detected in X-L digests of XanΔGL. These observations are consistent with the proposed role of GumF in selectively acetylating the internal Man residue rather than the terminal one.

Analysis of outer Man residues released from wild-type xanthan (WT Xan) revealed a greater diversity of modifications, including PMP-Man$^{Ac}$ (*m/z* 553), PMP-Man (*m/z* 511), PMP-Man$^{Pyr}$ (*m/z* 581) and doubly acetylated Man (*m/z* 595, observed as *m/z* 553 due to ionisation effects) (**Fig. S4**). Together, these results highlight the structural complexity of xanthan acetylation and provide a molecular basis for the variability in its biophysical properties, such as viscosity and solubility, observed across different production batches.

**Supplementary Result 2. *In vitro* production, identification, and stability assessment of short-chain acyl-CoA derivatives**

An LC-MS/MS-based method was used to analyse coenzyme A (CoA) and a series of short-chain acyl-CoA derivatives, primarily those containing acyl chains of two to four carbon atoms. These compounds were generated from both *in vitro* enzymatic reactions and intracellular metabolite extracts and were subsequently used as acyl donors in transferase assays.

Acyl-CoAs were detected by mass spectrometry operated in positive-ion mode. Protonated parent ions $[M+H]^+$ were observed at *m/z* 768.5, 836.0, 822.0, 810.0, 836.0 and 824.0, corresponding to CoA, crotonyl-CoA (CA-CoA), acryloyl-CoA (AA-CoA), acetyl-CoA (Ac-CoA), methacryloyl-CoA (MAA-CoA) and propionyl-CoA (PA-CoA), respectively (**Fig. S5a**). $MS^2$ spectra were acquired for all detected acyl-CoAs and provided characteristic fragmentation patterns that served as additional criteria for compound identification.

All protonated acyl-CoAs exhibited two dominant fragmentation pathways. The first involved cleavage between the pantothenate moiety and the β-phosphate of the adenosine group, generating an acyl–thioester–pantothenic acid fragment. The second pathway involved cleavage between the α- and β-phosphates, producing an adenosine monophosphate ion together with phosphate-derived fragments, including a diagnostic ion at *m/z* 428. These fragmentation behaviours are consistent with those previously reported by Perera et al. [4]. The first fragmentation pathway yielded acyl-CoA-specific ions, including *m/z* 329.1 for CA-CoA, 315.1 for AA-CoA, 303.0 for Ac-CoA, 329.2 for MAA-CoA and 317.1 for PA-CoA (**Fig. S5b**).

In reactions generating AA-CoA, an additional ion at *m/z* 840.0 accumulated over time, consistent with incorporation of a water molecule. The appearance of a corresponding fragment ion at *m/z* 333 (315 + 18) further supports this interpretation, suggesting hydration at the highly reactive carbon–carbon double bond of the acryloyl moiety.

Given the inherent instability of CoA thioesters—which can undergo degradation during the reaction, thereby reducing the effective concentration of the acyl donor in transferase assays and potentially falling below the detection limit—a time-course analysis was performed to evaluate the stability of CoA thioesters during Stlig-catalysed CoA production. In addition, spiking experiments were conducted during metabolite analysis to assess the stability and recovery of CoA species.

In the absence of authentic standards, relative quantification of individual acyl-CoA species at different time points was performed using Bruker Quant Data Analysis software. For each acyl-CoA, the most intense peak—validated by its *m/z* value and characteristic fragmentation pattern—was normalised to 100%, and relative abundances in other samples were calculated accordingly.

As shown in **Fig. S6**, MAA-CoA exhibited relatively high stability, retaining approximately 90% of its maximum abundance after 24 h, corresponding to the duration of the transferase assay. By contrast, Ac-CoA, PA-CoA and crotonyl-CoA (CA-CoA) retained approximately 60%, 20% and 80% of their initial abundances, respectively (**Fig. S6b–d**). Notably, AA-CoA underwent complete hydration within 10 min, consistent with its high chemical reactivity. To compensate for the intrinsic instability of acyl-CoAs and to ensure sufficient substrate availability during transferase assays, higher concentrations of Ac-CoA and PA-CoA were used. In addition, commercially available AA-CoA was employed in substrate promiscuity assays.

Following the introduction of the *Stlig* gene into *Xcc ΔgumL*, it was anticipated that supplementation with short-chain organic acids would lead to intracellular formation and detection of the corresponding CoA thioesters. However, neither CoA nor acyl-CoA species were detected in quenched cellular metabolite extracts. Spiking experiments, in which defined amounts of CoA or acetyl-CoA (100 µM) were added either before or after sample processing steps—including solvent evaporation and filtration—resulted in complete degradation of the compounds. These observations underscore the intrinsic instability of CoA thioesters and highlight the technical challenges associated with their detection in *Xcc* metabolite extracts.

**Supplementary Result 3. Short-chain organic acids utilization in *Xcc* for *in vivo* xanthan modification**

To achieve concentrations compatible with the detection limits of the in-house analytical methods, including HPLC and LC–MS/MS, for assessing *in vivo* catalytic promiscuity, the growth of *Xcc ΔgumL* (*pStlig*) was evaluated in LB medium supplemented with increasing concentrations of organic acids, including acrylic, propionic, crotonic and methacrylic acids, with the pH adjusted to 7.0. Elevated concentrations of short-chain organic acids, particularly acrylic acid, have been reported to inhibit bacterial growth by interfering with the fatty acid β-oxidation pathway, which may consequently reduce polysaccharide biosynthesis [5]. The *Xcc ΔgumL (pStlig)* strain was selected because it retains both GumF and GumG acetyltransferases (ACTases) while expressing *Stlig* from a plasmid. It was hypothesised that the supplemented organic acids could be converted into their corresponding CoA thioesters and subsequently incorporated into the polymer by one or both ACTases. Such incorporation would increase the degree of acylation and, in the case of acrylic acid, potentially enable the biosynthesis of a UV-curable acrylated polymer.

In this study, precultures were prepared, and harvested cell pellets were washed three times before inoculation into fresh LB medium supplemented with gentamicin, 1 mM IPTG and varying concentrations of the respective organic acids at an initial $OD_{600}$ of 0.1. Acrylic acid concentrations in the culture medium were quantified at the beginning and end of the growth period by HPLC using a Rezex™ ROA $H^+$ column. This analysis was performed to monitor acid assimilation, evaluate whether the supplied concentrations were sufficient to support bio-based transformation, and determine whether the resulting levels approached the detection limits of the in-house analytical methods used for polymer analysis.

As shown in **Fig. S7a**, supplementation with 10 mM acrylic acid significantly inhibited cell growth. However, the intracellularly assimilated acrylic acid concentration plateaued at approximately 5 mM, suggesting the presence of an uptake threshold (**Fig. S7b**). Therefore, 5 mM acrylic acid was selected as the feeding concentration for subsequent experiments, as this level remained compatible with the detection limits of the in-house HPLC and LC–MS/MS methods used for polymer characterisation. Other organic acids were supplemented at 10 mM, as no significant inhibitory effects on cell growth were observed at this concentration (**Fig. S7b**).

**Supplementary Result 4. Chemical stability and detection of acryloyl functional groups in free and CoA-Conjugated forms**

Given the high reactivity of the acryloyl group in acrylic acid and the harsh hydrolytic conditions used during sample preparation (90 °C for 16 h in 250 mM $H_2SO_4$), detection of acryloyl modifications on the polymer proved challenging, even if acyltransferase activity successfully mediated transfer of the group onto the polymer backbone. This limitation was evident from analysis of an acrylic acid standard using a Rezex™ ROA $H^+$ column with 2.5 mM $H_2SO_4$ as the mobile phase, which revealed two distinct peaks: a major peak (Peak 1) co-eluting with propionic acid (retention time, 21.69 min) and a minor peak (Peak 2) corresponding to intact acrylic acid (retention time, 23.25 min). These observations indicate that partial chemical conversion of acrylic acid, likely involving reduction or hydration of the α, β-unsaturated carbon–carbon double bond, may occur during chromatographic separation (**Fig. S8**).

This chemical conversion was further corroborated by forced hydrolysis experiments, in which wild-type xanthan spiked with 5 mM acrylic acid was incubated in 250 mM $H_2SO_4$ at 90 °C for 16 h. Under these conditions, acrylic acid was completely converted to propionic acid (**Fig. S8**), highlighting its pronounced susceptibility to acid-catalysed reduction. Collectively, these findings underscore a critical limitation of the in-house HPLC workflow: detection of propionic acid in xanthan samples derived from acrylic acid-supplemented *Xcc ΔgumL (pStlig)* cultures cannot unambiguously distinguish whether reduction of the C=C bond occurred *in vivo* during biotransformation or was introduced during post-hydrolysis sample processing.

**Supplementary Result 5. *In vivo* assay of GumG reveals catalytic promiscuity towards diverse acyl donors**

To further confirm that the propionyl group was specifically transferred to the outer mannose (Man) residue of xanthan by GumG upon feeding *Xcc ΔgumL (pStlig)* cells with propionic acid, an LC–MS/MS-based analysis was performed. This approach was required because the HPLC-based assay quantifies total propionic acid released from the polymer but does not discriminate between modifications catalysed by GumF or GumG (**Fig. 1e**). To localise the modification site, outer Man residues were enzymatically released from the polymer using xanthan lyase (X-L), and the resulting monosaccharides were purified as described in the Materials and Methods. As a negative control, xanthan produced by *Xcc ΔgumL* lacking *Stlig* expression and cultured in the presence of 10 mM propionic acid was analysed in parallel.

Extracted ion chromatograms revealed a prominent peak at *m/z* 567 in the experimental sample, corresponding to propionylated outer Man and supporting GumG-mediated transfer of the propionyl group (**Fig. S9a**). A minor peak at the same *m/z* value was also detected in the control sample, suggesting low-level background modification, potentially arising from catalytic promiscuity of endogenous CoA-dependent enzymes in *Xcc* (**Fig. S9a**).

Further validation was obtained by $MS^2$ fragmentation analysis. In contrast to unmodified Man, which exhibited a characteristic fragment ion at *m/z* 493.1, the propionylated outer Man lacked this signal and instead displayed a diagnostic fragment at *m/z* 549.1 (+56 Da), consistent with incorporation of a propionyl group (**Fig. S9b**). This fragment was detected exclusively in the experimental sample, confirming site-specific propionylation of the outer Man residue.

**Supplementary Result 6. Pseudo-acceptor xanthan pentamers as model substrates**

The activity of WT GumG toward pseudo-acceptor xanthan pentameric units (PenΔFGL and PenΔGL) was evaluated, and acetylated products were detected in the reaction mixtures but not in negative controls, as described above. To identify the specific acetylation sites catalysed by GumG, $MS^2$ fragmentation patterns of the acetylated products were analysed in detail. Using a PMP derivatisation strategy, the open-ring aldehyde at the reducing end of the sugar reacts with two PMP molecules to form a bis-PMP derivative. Consequently, only the reducing-end glucose (Glc) residue of the pentamer can be derivatised, enabling unambiguous localisation of acetylation sites [6].

When PenΔFGL was used as the acceptor substrate, characteristic fragment ions at *m/z* 553 (acetylated monomer), 715.3 (acetylated dimer), 877.4 (acetylated trimer), 1053.5 (acetylated tetramer) and 1217.5 (acetylated pentamer) were detected, consistent with acetylation of the reducing-end Glc residue (**Fig. S10b**). In addition, fragment ions corresponding to a non-acetylated monomer (*m/z* 511.2) together with acetylated higher-order fragments (715.3, 877.4, 1053.5 and 1217.5) were also observed, suggesting that acetylation can additionally occur on the second Glc residue from the reducing end (**Fig. S10b**). Notably, seven chromatographically distinct peaks corresponding to *m/z* 1217.5 were detected, indicating the presence of multiple positional isomers of singly acetylated pentamers (**Fig. S10a**). Differences in retention time are consistent with acetylation at different hydroxyl positions on the same Glc residue, resulting in altered hydrophobicity.

Further analysis of reactions containing PenΔFGL revealed a doubly acetylated pentamer (*m/z* 1257), consistent with previous observations reported by Stankowski et al [6]. $MS^2$ fragmentation produced a dominant ion at *m/z* 595, confirming double acetylation on the reducing-end Glc residue (**Fig. S10c**). In rare cases, dual acetylation on two separate Glc residues was also detected, supported by fragment ions at *m/z* 553 and 757 derived from the *m/z* 1257 precursor. Together, these observations reflect substantial structural heterogeneity in GumG-mediated acetylation patterns.

$MS^2$ fragmentation analysis of reactions using PenΔGL as the acceptor similarly identified the reducing-end Glc as the primary acetylation site, consistent with the results obtained for PenΔFGL. However, acetylation in the PenΔGL reaction was not restricted to a single position. Instead, a broader distribution of acetylation patterns—including positional variability and instances of double acetylation—was observed, comparable to that detected for PenΔFGL. Precise assignment of individual acetylated products to specific chromatographic peaks was limited by the absence of authentic standards and the inherent structural complexity of the pentameric substrates.

#### Supplementary Tables

**Supplementary Table 1.** Primers used in this study.

**Supplementary Table 2. Conservation analysis of GumG residues based on ConSurf.** Conservation scores for 356 amino acid residues of GumG were calculated from a multiple sequence alignment of 49 homologous proteins using ConSurf.
